# Extending intravitreal retention of CNTF via fusion to the HA-binding protein VG1

**DOI:** 10.64898/2026.11.09.765452

**Authors:** Riley A. Suhar, Xiaowu Gu, Alexandra Robles, Robert F. Kelley, Devin B. Tesar

## Abstract

Glaucoma is the leading cause of permanent blindness worldwide. Vision loss in glaucoma stems from the loss of retinal ganglion cells (RGC). Ciliary neurotrophic factor (CNTF) has been shown to protect RGCs in pre-clinical models of glaucoma. However, CNTF has a short ocular retention time following intraocular injections due to rapid diffusion. To address this, we developed a fusion protein between CNTF and the hyaluronic acid-(HA) binding G1 domain of endogenous extracellular matrix (ECM) protein, versican (VG1; CNTF-VG1). We hypothesize that a CNTF-VG1 protein will have an extended ocular retention time compared to CNTF due to HA-binding with endogenous vitreous-HA. We demonstrate that CNTF-VG1 non-covalently binds HA with high affinity and is bioactive. We show that the vitreous and thermal stability of CNTF-VG1 can be significantly increased through co-formulation with HA. In an *in vivo* pharmacodynamic study, we show that VG1 significantly increases the window of neuroprotective JAK/STAT3 pathway activation in un-injured mice (C57BL/6J, female) retina. Finally, we close our work by showing that CNTF is neuroprotective of primary mouse cortical neurons in an acute H_2_O_2_ cytotoxicity model. Taken together, our work demonstrates the potential of HA-binding domain VG1 to improve the ocular retention of neuroprotective CNTF following intraocular injection.

## INTRODUCTION

Glaucoma is the leading cause of permanent blindness worldwide.^1^ One study tabulated that direct medical costs alone totaled approximately $2.9 billion in the US market.^2^ However, this is likely an underestimate, because as many as 50% of glaucoma patients go undiagnosed due to its asymptomatic presentation in its early stages.^3,4^ Despite these major patient and financial burdens, there is no cure for glaucoma and therapeutic options remain preventative at best. Elevated intraocular pressure (IOP) has been strongly implicated in glaucomatous progression,^5–7^ and it is thought to trigger a complex feedback loop ultimately leading to progressive damage and loss of the retina’s sole output neurons, the retinal ganglion cells (RGCs).^8^ Because of this, currently available therapies aim to specifically reduce elevated IOP, either through administration of eye drops^9,10^ or surgical intervention.^10,11^ However, IOP is not diagnostic. There are patients with healthy IOP and progressive neuropathy (“normal tension glaucoma”), patients with elevated IOP and no vision loss (“glaucomic suspects”); and importantly, decreasing IOP does not guarantee prevention of vision loss.^12^

While IOP is known to contribute to progressive blindness, it may only be part of the disease pathology. There is a growing body of evidence that RGC loss may be tied to a milieu of other physical, molecular, and chemical assaults. In live glaucomatous patient cohorts, for example, there is evidence of: reduced ocular blood flow,^13,14^ heightened mitochondrial dysfunction,^15^ and reduced neurotrophic factor concentration^16,17^ compared to healthy patients. Many of these findings have been recapitulated in pre-clinical animal models of glaucoma: tissue stiffening and fibrosis,^18^ mitochondrial dysfunction in RGC,^19^ glial cell activation,^20,21^ apoptosis-induced RGC death^22^ and evidence of heightened reactive oxygen species-(ROS) induced damage^23^ have all been recapitulated in animal models of glaucoma. Genetically, things do not look clearer; a recent genome-wide association study (GWAS) identified as many as 312 independent risk-loci for heritable primary open-angle glaucoma (POAG).^24^ Because of this complexity, the current paradigm is neuroprotection: instead of treating the numerous potential causes, we instead develop therapies that make RGCs more resistant.

Neurotrophic factors are highly potent cytokines that enact pro-survival machinery in neurons and have the potential to rescue dying RGCs to prevent further degradation. Brain-derived neurotrophic factor (BDNF),^25,26^ glial cell-line derived nerve growth factor (GDNF),^27^ nerve growth factor (NGF),^28,29^ and ciliary neurotrophic factor (CNTF)^30–32^ have all shown evidence of efficacy in pre-clinical models of glaucoma. Among these, CNTF is by far the most studied due to its high potency and activation of multiple pro-survival pathways (*e*.*g*., JAK/STAT, MAPK/ERK, and PI3K/AKT). However, a significant barrier to clinical translation of CNTF is maintaining a high enough retinal bioavailability. Post-injection, CNTF rapidly clears from the injection site with a plasma half-life of just 2 ∼ 6 hours in systemic administration.^33^ Additionally, because there is generally low retinal bioavailability in the retina following systemic^34^ and topical^35^ application of therapeutics, coupled with the potential for dose-dependent off-target side-effects,^33^ intravitreal (IVT) injection of CNTF is required to maximize retinal bioavailability. However, due to its rapid diffusion, this may necessitate repeat. *Neurotech* has proposed circumnavigating this challenge through a cell-based ocular implant (NT-501) to offer continuous CNTF delivery to the retina which has shown good safety tolerance in Phase 1 trials.^36,37^ However, not all patients are good candidates for surgical implantation, and alternative modalities of delivery may further increase accessibility of ocular therapies.

To address this, we investigated the hyaluronic acid-(HA) binding G1 domain of extracellular-matrix (ECM) protein versican (VG1) as a potential long-acting delivery (LAD) strategy for CNTF following IVT injection. The vitreous is composed of a gel-like substance that is rich in HA and collagen. VG1 non-covalently binds to HA through the formation of a link-protein complex.^38,39^ Therefore, we hypothesize that a fusion protein between VG1 and CNTF (CNTF-VG1) will have an enhanced window of neuroprotection compared to wild-type (WT) CNTF through a combination of reduced diffusion and immobilization to endogenous HA (Figure 1). Our lab recently showed that a VG1-Fab fusion has increased brain-retention following intracerebroventricular (ICV) administration,^40^ and another study showed a 3∼4 fold increase in ocular retention in rabbit and monkey eyes using a different HA-binding moiety, TSG-6, for an anti-VEGF antibody following IVT injection.^41^ The same moiety that we propose, VG1, was previously proposed as an LAD for IGF-1 in articular cartilage repair, however the available results were limited to *in vitro* characterization.^42^ To our knowledge, this is the first manuscript investigating the use of VG1 as a potential LAD technology for extended delivery of neurotrophic factors to the eye. In this study, we explore the feasibility of using HA-binding domain VG1 as an LAD for extending the ocular retention of CNTF following IVT injection. Taken together, our study offers a thorough characterization of HA-binding *via* fusion with VG1 as a potential LAD technology for CNTF delivery to the eye. More importantly, it offers a perspective on the benefits and challenges of using ECM-binding as a strategy for LAD.

**Figure 1.**
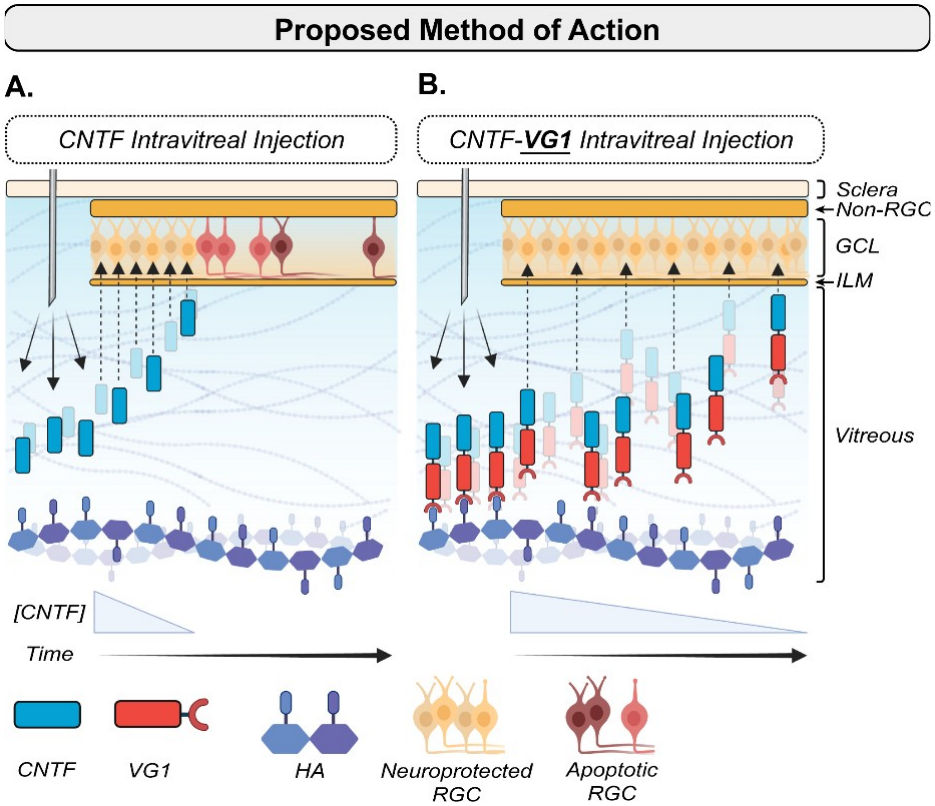
*Proposed method of action*. Simplified schematic comparing (A) wild-type CNTF and (B) CNTF-VG1 post-injection diffusion timelines. We hypothesize that a fusion protein between ciliary neurotrophic factor (CNTF) and the hyaluronic acid (HA)-binding domain G1 from versican (VG1; CNTF-VG1) will have an extended neuroprotective window compared to wild type CNTF due to the molecules reduced diffusion away from the injection site into the retina through non-covalent interactions with endogenous HA. Created in *BioRender*. Suhar, R. (2026) https://BioRender.com/xcxult6.

## MATERIALS AND METHODS

### Construct expression and purification in Chinese hamster ovary (CHO) cells

Constructs consisting of amino acids for: 1) recombinant human CNTF with a C-terminal histidine-6 (his6) tag (CNTF; ∼23.8 kDa), 2) Leu21 - Lys349 of mature, human, ECM protein versican with a C-terminal his6 tag (VG1; ∼36.3 kDa), 3) a fusion protein of CNTF with an N-terminal VG1 and C-terminal his6 tag (CNTF-VG1; ∼60.1 kDa), and 4) a non HA-binding CNTF-VG1 variant with a double-point (D197A, Y208A) mutation in the VG1 domain, CNTF-VG1^(ΔD197A,ΔY208A)^ with a C-terminal his6 tag (CNTF-ΔVG1; ∼60.1 kDa) were synthesized (Genscript, Piscataway, NJ) and expressed in Chinese hamster ovary (CHO) cells. Target proteins were purified using Ni-NTA column chromatography. Samples were fractionated to a purity of 95%+ monomer and stored long term at 4 °C in a storage buffer (20mM Histidine Acetate, 0.15M NaCl, pH5.5) until use.

### Coomassie and Western blotting for construct validation

Denatured purified proteins were separated on a 4 - 12% Bis-Tris pre-cast gel (ThermoFisher) in 1X MOPS SDS Buffer (NP0001, ThermoFisher). Separated gels were stained in SimplyBlue (LC6065, Invitrogen) for 1 hr. at room temperature, washed, and imaged (ChemiDoc, BioRad) for Coomassie blue visualization. For Western blotting, gels were dry transferred to a nitrocellulose (NC) membrane (Invitrogen) using an iBlot2 system, blocked in SuperBlock buffer (37516, ThermoFisher), and immunoblotted for his6 (1:1000; 2365S, CellSignaling) using LICOR Odyssey imager. Molecular weight (M_w_) range was determined using a standard ladder (LC5925, ThermFisher).

### Size exclusion chromatography (SEC) and fractionation

Purified proteins were analyzed by size exclusion chromatography (SEC) using a TSKgel SuperSW2000 (0018674, TOSOH Bioscience) connected to a high-performance liquid chromatography system (HPLC; Agilent 1290 Infinity). The column was equilibrated with mobile phase (0.2M K_3_PO_4_, 0.25 M KCl, 10% isopropanol, pH 6.2, isocratic, 30 min.) at room temperature, and 20 ∼ 100 µg protein was injected into the column and eluted at a flow rate of 0.5 mL min^-1^. The elutant was monitored by UV-Vis (λ = 280 nm), and representative traces were visualized and quantified using Chromeleon (ThermoFisher). If samples were < 95% monomeric, aliquots were loaded onto Superdex™ 10/300 GL column (17517601, Cytiva) equilibrated with storage buffer (20 mM Histidine Acetate, 0.15 M NaCl, pH5.5) and fractionated using the ÄKTA Pure system (Cytiva). The monomeric peak was collected and concentrated as needed using a spin column (10 kDa MWCO, Amicron).

### Hyaluronic acid (HA) binding kinetics evaluation using surface plasmon resonance (SPR)

Binding kinetics were measured using a Biacore T200 system (Cytiva). Biotinylated hyaluronic acid (HA) (10 kDa; HA-Mono-Biotin-10k, HAWorks) was immobilized on a streptavidin-functionalized sensor chip (BR100398, Cytivia) using EDC-NHS coupling in HBS-P buffer (BR100671, Cytiva). The test analyte was prepared in a 2X dilution series (5 µM ∼ 2.5 nM; HBS-P) and flowed over the HA-functionalized chip at a rate of 40 µL min^-1^. Association and dissociation phases were monitored for 180s and 120s, respectively. Chips were then regenerated using 4.0 M magnesium chloride hexahydrate (HR2-803, Hampton Research). Data were analyzed using the Biacore Evaluation Software to determine relevant binding kinetics: association rate constant (*k*_*a*_), dissociation rate constant (*k*_*d*_), and equilibrium dissociation rate (*K*_*D*_).

### TF-1 proliferation assay for measuring construct relative bioactivity

Human myeloid leukemia suspension cells (TF-1) were grown in complete medium (RPMI 1640, 2 mM L-glutamine, 1 mM sodium pyruvate, 2.5 ng mL^-1^ GM-CSF (215-GM, R&D Systems), 10% fetal bovine serum (FBS)). For the proliferation assay, cells were collected *via* centrifugation and carefully washed 3 times in pre-warmed GM-CSF-free medium, filtered using a 40-µm cell strainer (CLS431750, Millipore Sigma) and measured for viability and cell density using trypan blue and an automated cell counter (Countess 3, ThermoFisher). Cells were then re-suspended in testing medium (GM-CSF-free, 0.25 µg mL^-1^ CNTFRα (303-CR-050/CF, R&D Systems) and incubated for ∼10 min. Constructs were prepared at 10X concentration in testing medium over a 9-point, 5-decade concentration range (200 nM - 0.002 nM) with half-logarithmic intervals and an additional construct-free (testing medium only) negative control (10 total conditions). 10 µL of 10X construct and 90 µL of TF-1 cells were added to a final concentration of 20,000 cells/well in ultra-low attachment 96-well tissue culture plates (3474, Corning), in triplicate, and incubated for 72 hours. Cell proliferation was measured on a plate reader using an MTS colorimetric assay (G3582, Promega), per manufacturer instructions. ED_50_ was interpolated using a sigmoidal, 4PL curve fit in PRISM (GraphPad).

### CNTF-VG1 thermal and low-HA solution stability screen

CNTF-VG1 was buffer-exchanged into PBS using a desalting column (89882, ThermoFisher) and mixed with solutions of HA (LifeCore) to a final [CNTF-VG1] of 5 mg mL^-1^. HA was varied for M_w_ (10, 20, 40, 60, and 100 kDa) and concentration

(50, 500, 1250, 2500, 5000, 10,000 15,000, and 20,000 µg mL^-1^). CNTF-VG1 + HA solutions, alongside PBS-only and CNTF-VG1-only (no HA), were imaged in bright field microscopy for qualitative assessment of precipitation after 24 hours of incubation at room temperature and subsequently after 24 hours of incubation at 37 °C.

### HA co-formulation effect on vitreous compatibility using background membrane imaging (BMI)

CNTF-VG1 was buffer-exchanged into PBS using a desalting column (89882, ThermoFisher). For the HA co-formulation group (CNTF-VG1 + HA), CNTF-VG1 was carefully mixed with 60 kDa HA (LifeCore) using gentle pipetting to a final [CNTF-VG1] of 5 mg mL^-1^ and [HA] of 1% (w/v) and then left to incubate overnight at room temperature with gentle agitation on an automatic rotator. The following day, pre-chilled (4 °C) PBS, CNTF-VG1 and CNTF-VG1 + HA were pipetted at a 1:3 ratio into pre-warmed (37 °C) PBS, isolated pig vitreous (BioIVT), and primary rat vitreous (in triplicate) and left to incubate overnight at 37 °C. The following day, each injection condition was carefully mixed to form an even suspension, and aliquots from each were further diluted 10X in pre-warmed PBS. 40 µL of diluted sample was then carefully pipetted onto a 96-well format, 0.4 µm pore filter membrane plate (SolveScientific) and the precipitates were immobilized onto the optically transparent membrane using an in-line vacuum. Each sample was washed 3 times with ultra deionized water to remove salt deposits, imaged (4X Brightfield) and precipitation was quantified using an Aura PTx background membrane imaging system (Halo Labs).

### Intravitreal injection in uninjured mice eyes and retina dissection

All animal care and handling procedures were reviewed and approved by the Genentech Institutional Animal Care and Use Committee (IACUC) and were conducted in full compliance with regulatory statutes, IACUC policies and in accordance with the ARVO statement for the Use of Animals in Ophthalmic and Vision Research. All animals were housed in a specific pathogen-free facility and on a 12h light/12h dark cycle with ad libitum access to food and water, and monitored daily. Female, 6–8-week-old C57BL/6J mice (stock #000664, Jackson Lab) were used. On Day 0, mice were deeply anesthetized and eyes were dilated with 1% tropicamide drops. Under a surgical microscope, a 33G 1/2 TSK STERiJECT hypodermic needle was used to puncture the posterior limbus. A 34G blunt Hamilton syringe was then used to deliver a 1.5 µL injection of 2 µg µL^-1^ CNTF (total: 3 µg, or molar equivalent CNTF-VG1) at an approximate rate 1 µL min^-1^, by hand. Following IVT, a topical antibiotic (Neomycin and polymyxin B Sulfate and Bacitracin Zinc Ophthalmic Ointment; Bausch & Lomb) was applied, and an extended-release analgesic (3.25 mg/kg Ethiqa XR; Fidelis Pharmaceuticals) was administered. Animals were placed on a pre-warmed heating pad at 37°C for recovery. At target time points, mice were sacrificed with carbon dioxide asphyxiation followed by cervical dislocation retinas were carefully dissected, snap-frozen in liquid nitrogen, and stored at -80 °C until processing. Treatment groups were not blinded and were randomly assigned at day 0; any mice with evidence of ocular inflammation or stress were sacrificed per veterinarian guidance and excluded from quantification. All mice were stored in the same housing location, and retinas were collected from sacrificed mice in the order in which they were injected to minimize errors in measurement.

### pSTAT3/STAT3 quantification in isolated retina via immunoblotting

Retinas were lysed in RIPA buffer (9806, Cell Signaling) with protease (1861279, ThermoFisher) and phosphatase inhibitor (78427, ThermoFisher) inhibitor. Protein content was measured using a bicinchoninic acid (BCA) assay (23225, ThermoFisher). ∼20 µg of tissue lysate was separated *via* electrophoresis, transferred to a NC membrane, and blocked as previously described. Immunoblotting and imagining were conducted in two stages: first, membranes were co-labeled with antibodies for pSTAT3^Tyr705^ (1:1000, Rb; 9145, Cell Signaling) and β-Actin (1:10,000, ms; 3700, Cell Signaling) overnight (ON) at 4 °C, washed, and subsequently stained with a corresponding secondary antibody cocktail (1:10,000, anti-Rb; LIC-926-3221, 1:20,000 anti-ms; LIC-926-68052, LiCOR) for 1 hr. at room temperature, washed, and imaged (LiCOR Odyssey). Membranes were stripped (1 hr., room temperature; 21059, ThermoFisher), washed, re-blocked and then incubated ON at 4 °C with antibodies for STAT3 (1:1000, Rb; 12640, Cell Signaling) and β-Actin, washed and subsequently stained with a corresponding secondary antibody cocktail for 1 hr. at room temperature, washed and imaged (LiCOR Odyssey). For quantification, band intensities were measured in Fiji^43^ (ImageJ 1.54 p) with the Gel Analyzer tool. Band intensities for pSTAT3^Tyr705^ and STAT3 were first normalized to β-Actin and then quantified for each condition and time point as a ratio of pSTAT3^Tyr705^/STAT3. For the preliminary day 1 and day 7 experiment, two individual retinas were averaged from two separate mice treated with either CNTF, CNTF-VG1, and CNTF-VG1 + HA or vehicle only (PBS; control) at 1- and 7-days post IVT injection; a naive (untreated) retina was additionally shown for qualitative comparison but not used in quantification. For the time-course experiment, retinas were bilaterally injected with either CNTF-VG1, and CNTF-ΔVG1 or CNTF-VG1 + HA and collected 1-, 3-, 5-, and 7-days post IVT injection. For quantification, both retinas were averaged from each animal, and 3 animals were used in each group at each time point. In total, 56 mice were used in analysis.

### Primary mouse cortical neuron (MCN) isolation and culturing

All animal care and handling procedures were reviewed and approved by the Genentech Institutional Animal Care and Use Committee (IACUC) and were conducted in full compliance with regulatory statutes, IACUC policies and in accordance with the ARVO statement for the Use of Animals in Ophthalmic and Vision Research. E15 ∼ E15.5 pregnant C57BL-6N mice (Charles River, strain number: 027) were sacrificed with carbon dioxide asphyxiation followed by cervical dislocation. The uterine sac was excised, embryos were removed, and the brains were carefully dissected and stripped of all meninges. Tissue was carefully washed with successive steps by moving progressively cleaned tissue to fresh, pre-chilled (4 °C) HBSS (14170120, Gibco). The cortices were isolated from the central cortex, olfactory bulb, and hippocampus and then collected for dissociation. Cortices were washed (3X) with HBSS^-/-^, dissociated with 0.25% trypsin in HBSS^-/-^ + DNAse I (10 min, 37 °C); trypsinization was neutralized with pre-chilled plating medium (Neurobasal, 20% heat-inactivated horse serum (26050-088, Invitrogen), 1X GlutaMAX, and 25 mM sucrose) and further washed (3 times) with pre-chilled HBSS^-/-^ before being re-suspended in plating medium. Cortices were homogenized with gentle trituration, centrifuged, re-suspended in plating medium, filtered using a 40-µm cell strainer (CLS431750, Millipore Sigma) and measured for viability and cell density using trypan blue and an automated cell counter (Countess 3, ThermoFisher). Finally, cells were seeded at approximately 1.25 M cm^-2^ (*e*.*g*., ∼40,000 cells / well in a 96-well plate) in a Poly-L-Lysine-coated plate and left to incubate overnight in a CO_2_ humidity-controlled incubator. The following day, the plating medium was aspirated and replaced with fresh, pre-warmed NbActiv4 medium (NC1477957, FisherScientific) and left to culture for 5 ∼ 7 days prior to additional experimentation.

### CNTF neuroprotection assay against acute H_**2**_**O**_**2**_ exposure in mouse cortical neurons (MCN)

To assess the neuroprotection of CNTF against oxidative stress-induced cell death, we leveraged an H_2_O_2_ cell-cytotoxicity model. H_2_O_2_-treatment groups were divided into two subgroups: 1) CNTF-free or 2) CNTF pre-treated, where cells were cultured with 100 ng mL^-1^ CNTF overnight prior to addition of H_2_O_2_. H_2_O_2_ exposure was conducted either chronically or with acute exposure; in the acute condition, H_2_O_2_ was neutralized using catalase (C9322, Sigma) after either 15, 30, 45, or 60 minutes of exposure to attenuate cell damage and present a range of insults. For testing purposes, 5 - 7-day old cultures of primary MCN were divided into 4 groups (n = 5 wells / group): 1) untreated, 2) catalase pre-treated (catalase added 1 hr. prior to addition of H_2_O_2_), 3) chronic H_2_O_2_ exposure, and 4) acute H_2_O_2_ exposure. Both acute and chronic groups had two sub-groups: CNTF-free and CNTF pre-treated. Cell death was monitored with Cytotox Red (4632, Sartorius) every 6 hours over the course of 72 hours with live cell imaging (Incucyte, Sartorius), normalized to the highest average reported cell death on a given plate and used to estimate a percent cell-death for each time point. Results are presented as an average of 3 independent MCN preparations.

### Statistical analysis

These data were analyzed and graphed using Prism (GraphPad). For all measurements, a significance level of α = 0.05 was used and level of significance is represented as follows: * p < 0.05; ** p < 0.01; *** p < 0.001; **** p < 0.0001; ns = not significant in all cases. All data is presented as mean ± standard deviation (SD); where informative, individual replicates are included (•). The specific statistical test employed, and replicate count, is listed in the figure captions where appropriate; validation of the assumptions of the statistical methods employed were checked using built-in tools within Prism.

## RESULTS

### Fusion of CNTF with VG1 allows for high-affinity binding to HA without significantly affecting bioactivity of CNTF

The first aim of the study was to demonstrate that the HA-binding affinity, as well as bioactivity of CNTF, are conserved following fusion with VG1. For this analysis, we expressed three constructs in CHO cells: CNTF, CNTF-VG1, and VG1 (Construct schematics in Figure S1). As shown in Figure 2A,B, Coomassie and Western blots revealed high purity products that were monomeric (95%+) by SEC (Figure 2C). Monomeric, low molecular weight species (LMWS), and high molecular weight species (HMWS) are indicated within the figure.

**Figure 2.**
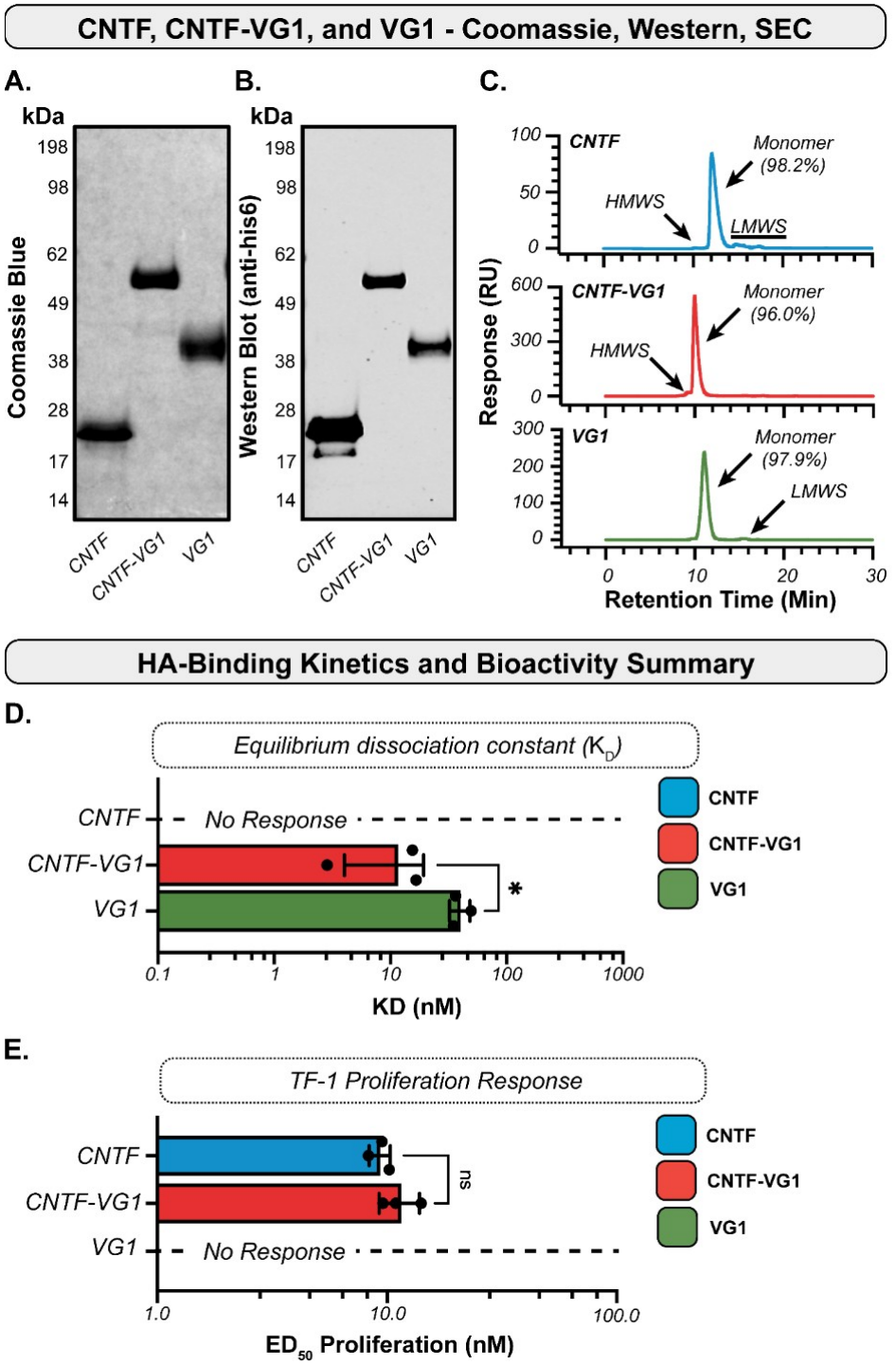
*Characterization of recombinant CNTF and CNTF-VG1 for HA-binding kinetics and relative bioactivity*. (A) Representative Coomassie stain (grayscale), (B) western blot (anti-his6) and (C) size exclusion chromatography (SEC) traces of CNTF, CNTF-VG1, and VG1 constructs. Monomeric, high molecular weight (HMWS) and low molecular weight species (LMWS) are labeled. (D) Tabulated equilibrium dissociation constant (*K*_*D*_) against immobilized hyaluronic acid in surface plasmon resonance. (E) Tabulated equilibrium ED_50_ of proliferation in human TF-1 proliferation cell-assay. Statistics: Unpaired two-tailed student’s T-test; Data presented as mean ± SD; α = 0.05; * p < 0.05; ns = not significant; n = 3 (•).

Following expression and purification, we measured the kinetic parameters of HA-binding using SPR against immobilized HA (Representative SPR curves: Figure S2A). Our measurements indicate that while CNTF did not have measurable non-covalent binding with HA, CNTF-VG1 bound to HA with high affinity with an equilibrium dissociation constant (*K*_*D*_) of ∼12 nM. Interestingly, this is a significantly higher (p = 0.0112; *) affinity than was measured for VG1 alone (Figure 2D; *K*_*D*_ ∼40 nM). Both molecules had approximately the same association constant (*k*_*a*_); however, CNTF-VG1 had a ∼3X lower dissociation constant (*k*_*d*_) potentially suggesting that, when immobilized, secondary, non-specific, interactions may be the principal contributor to the measured higher-affinity. A complete breakdown of kinetic parameters is summarized in Figure S2B.

To demonstrate that bioactivity of CNTF was not impacted by its fusion to VG1, we leveraged the IL-6-expressing human TF-1 cell line which has been shown in literature previously to respond proliferatively to CNTF.^44,45^ As shown in Figure 2E, there was no statistically significant difference in equilibrium ED_50_ of proliferation between CNTF and CNTF-VG1, suggesting that bioactivity was not impacted by the presence of VG1 (Representative curves shown in Figure S2C; Tabulated ED_50_ results in Figure S2D). To validate this in a more relevant cell-line, we further showed that both of our constructs, CNTF and CNTF-VG1, activate the JAK/STAT3 pathway in human embryonic stem-(ESC) derived RGC (hESC-RGC) (Figure S3A) and tabulate that there is no measurable difference in pSTAT3^Tyr705^ activation as measured by immunoblotting (Figure S3B). Lastly, in anticipation of *in vivo* ocular retention studies in mice models, we additionally validated cross-reactivity of our molecules in primary mouse cortical neuron (MCN) cultures and showed that human CNTF activates the JAK/STAT3 pathway at a similar level to mouse CNTF *in vitro* (Figure S3C-D). Taken together, these *in vitro* results show that fusion of VG1 to CNTF imparts HA-binding while not affecting its bioactivity.

### Co-formulation with HA significantly increase vitreous and thermal stability of CNTF-VG1

Our own lab has previously reported that fusion-antibodies with HA-binders such as TSG-6 and VG1 can experience significant precipitation when introduced to vitreous, and that these effects can be qualitatively ameliorated by pre-complexing with an excess of HA.^46^ To study this phenomenon further, and its effect on *in vivo* ocular retention, we developed a series of tests to quantify vitreous-induced precipitation *ex vivo*. First, to establish a relevant range of HA solutions, we measured [HA] in a variety of commercially available (human, pig, Cynomolgus (cyno) monkeys, rabbit) and (Sprague Dawley, rat) vitreous samples using an HA ELISA assay (Figure S4). Assuming [HA] is similar in rats and mice, and an approximate vitreous volume of 4.5 µL in mouse eyes,^47^ we estimate based on our measurements that mice have an HA content of approximately ∼ 7.2 ng / eye. Given that typical dosing ranges for CNTF in other mice studies range from 1 - 4 µg, for a molar equivalent of CNTF-VG1 this represents an approximate VG1:HA repeat-unit ratio of ∼2.4 - 9.5. In practice, this could potentially mean that a large fraction of our dose may not have the opportunity to bind with an endogenous HA partner due to the low vitreous HA content in mice vitreous.

To screen for vitreous compatibility challenges, we prepared solutions of CNTF-VG1 and synthetic HA (Lifecore), wherein the HA concentration was varied from ∼50 µg mL^-1^ to as high as 20,000 µg mL^-1^ for a range of commercially available HA’s (M_w_: 10 - 100 kDa) and a fixed final concentration of CNTF-VG1 (5.0 mg mL^-1^). Samples from each condition were qualitatively assessed with brightfield imaging after overnight incubation at room (23 °C) and physiological (37 °C) temperatures to study the effect of HA-binding and temperature on CNTF-VG1 solution stability. Our screen revealed three major findings, which we have summarized with a selection of images from our full array (Figure 3A-H). First, CNTF-VG1 is thermally unstable; after overnight incubation at 23 °C, we observed evidence of precipitation in the CNTF-VG1-only group (Figure 3B), which is exaggerated at 37 °C (Figure 3F). Second, we found that in our lowest HA concentration group (50 µg mL^-1^), there was significant precipitation compared to our CNTF-VG1-only group (Figure 3C,G), which is likewise exaggerated after overnight incubation at 37 °C. Finally, and most interestingly for our work here, we observed that both of these temperature and HA-binding induced precipitation events are completely ablated at sufficiently high HA concentrations (Figure 3 D,H), as indicated by no observable precipitation for samples where [HA] > ∼1000 µg mL^-1^

**Figure 3.**
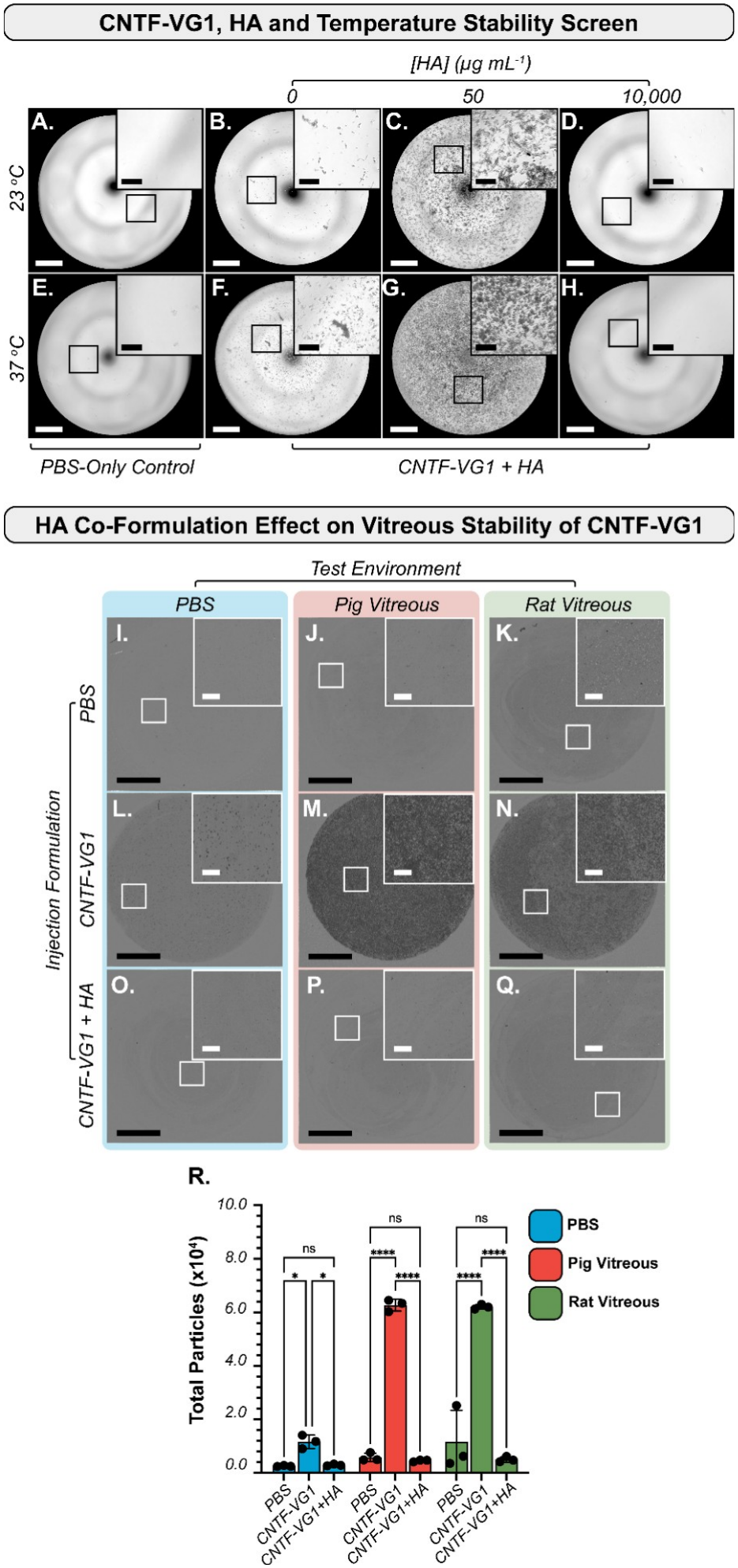
*CNTF-VG1 thermal and vitreous stability optimization*. (A - H) Representative bright field images from a stability screen of CNTF-VG1 (5.0 mg mL^-1^) and HA at room temperature and 37 °C. (I - Q) Representative background membrane imaging (BMI) images from HA-coformulation stability test quantifying precipitation of PBS, CNTF-VG1, and CNTF-VG1 + HA (1%, 60 kDa) following overnight incubation at 37 °C in PBS, pig vitreous, and rat vitreous. (R) Quantified precipitation using BMI. Statistics: One-way ANOVA, mixed model, multiple comparisons with post-hoc Tukey correction; data presented as mean ± SD; α = 0.05; * p < 0.05; **** p < 0.0001; ns = not significant; n = 3 (•).

(Figure 3D,G; Figure S5,6). This presents a potential two-in-one solution: co-formulation (pre-complexing) CNTF-VG1 with HA may both supplement the low vitreous HA content in eyes to maximize the amount of HA-bound CNTF while also improving thermal and vitreous stability. Images of our full array, and relevant controls, can be found in Figure S5 and Figure S6 for 23 °C and 37 °C conditions, respectively.

However, these findings led us to two additional questions. First, how does the diffusivity of CNTF-VG1 co-formulated with synthetic HA compared to its diffusivity when it is bound potentially higher M_w_ vitreous HA and, secondly, does co-formulation prevent precipitation of CNTF-VG1 in vitreous at physiological temperatures? To select a co-formulation condition to study these questions, we qualitatively assessed the injectability of our different CNTF-VG1:HA formulations and found that the highest [HA] and M_w_ formulation condition that we could inject through a 33G needle was 1% (w/v) 60 kDa HA (data not shown).

To evaluate the effect of pre-complexing on diffusion, we fluorescently tagged CNTF-VG1 with AF488 and measured its diffusivity (D; µm^2^ s^-1^) in highly dilute solutions (CNTF-VG1 << [HA]) of commercial 60 kDa HA, pig vitreous, rat vitreous, and PBS (negative control) using fluorescence correlation spectroscopy (FCS). Our results suggest solutions with HA have a nearly 25 - 50X decrease in diffusivity compared to PBS (D = 54.25 µm^2^ s^-1^), with measured diffusivity of 1.62 µm^2^ s^-1^ in pig vitreous, 0.904 µm^2^ s^-1^ in rat vitreous, and 2.35 µm^2^ s^-1^ in commercial 60 kDa HA (FCS data summarized in: Figure S7). Taken together, these results show that HA-binding both significantly reduces the diffusivity of CNTF-VG1, and that the reduction in diffusivity is similar between synthetic 60kDa HA and vitreous HA.

Finally, to evaluate whether HA co-formulation prevents precipitation of CNTF-VG1 in vitreous we designed an assay to measure thermal and vitreous-induced precipitation. In brief, solutions of PBS (positive control), CNTF-VG1, and CNTF-VG1 + HA were mixed with pre-warmed solutions of PBS, rat vitreous, and pig vitreous and incubated overnight at 37 °C to emulate IVT conditions. We then leveraged background membrane imaging (BMI) to approximate the number of precipitates in each sample to quantify the effect of formulation on protein solubility. A schematized graphic of this protocol is available in Figure S8 and example particle masking images generated in BMI from samples of low- and high-precipitation samples can be found in Figure S9. The results reveal a clear difference in precipitation between PBS, CNTF-VG1, and CNTF-VG1 + HA groups (Representative BMI images: Figure 3I-Q), with CNTF-VG1 having significantly higher precipitation events in all conditions than either PBS or CNTF-VG1 + HA across all conditions (Figure 3R). Taken together, these data suggest that co-formulating CNTF-VG1 with HA may confer increased thermal and vitreous stability while maintaining comparable reduction in diffusivity as binding to endogenous HA post-IVT injection.

### VG1 significantly increases JAK/STAT3 activation window of CNTF in mouse retina following IVT injection

To study the effect of VG1 and HA co-formulation on ocular retention, we conducted a pharmacodynamic (PD) study by measuring the relative activation of the CNTF downstream neuroprotective signaling pathway, JAK/STAT3, following IVT injection, in mice, retina. In brief, a bilateral IVT injection was administered in un-injured C57BL/6J (♀) mice (containing either vehicle (PBS), CNTF, CNTF-VG1, or CNTF-VG1 + HA. We then isolated the retinas and quantified the relative pSTAT3^Tyr705^ over total STAT3 protein intensity with immunoblotting at 1 and 7-day post-administration. A schematic summary of our study design is in Figure 4A, and representative blots along with tabulated results can be found in Figure 4B and Figure 4C-D, respectively. We derived a few key findings from our preliminary study. First, elevated JAK/STAT3 signaling in retinas dosed with CNTF and

**Figure 4.**
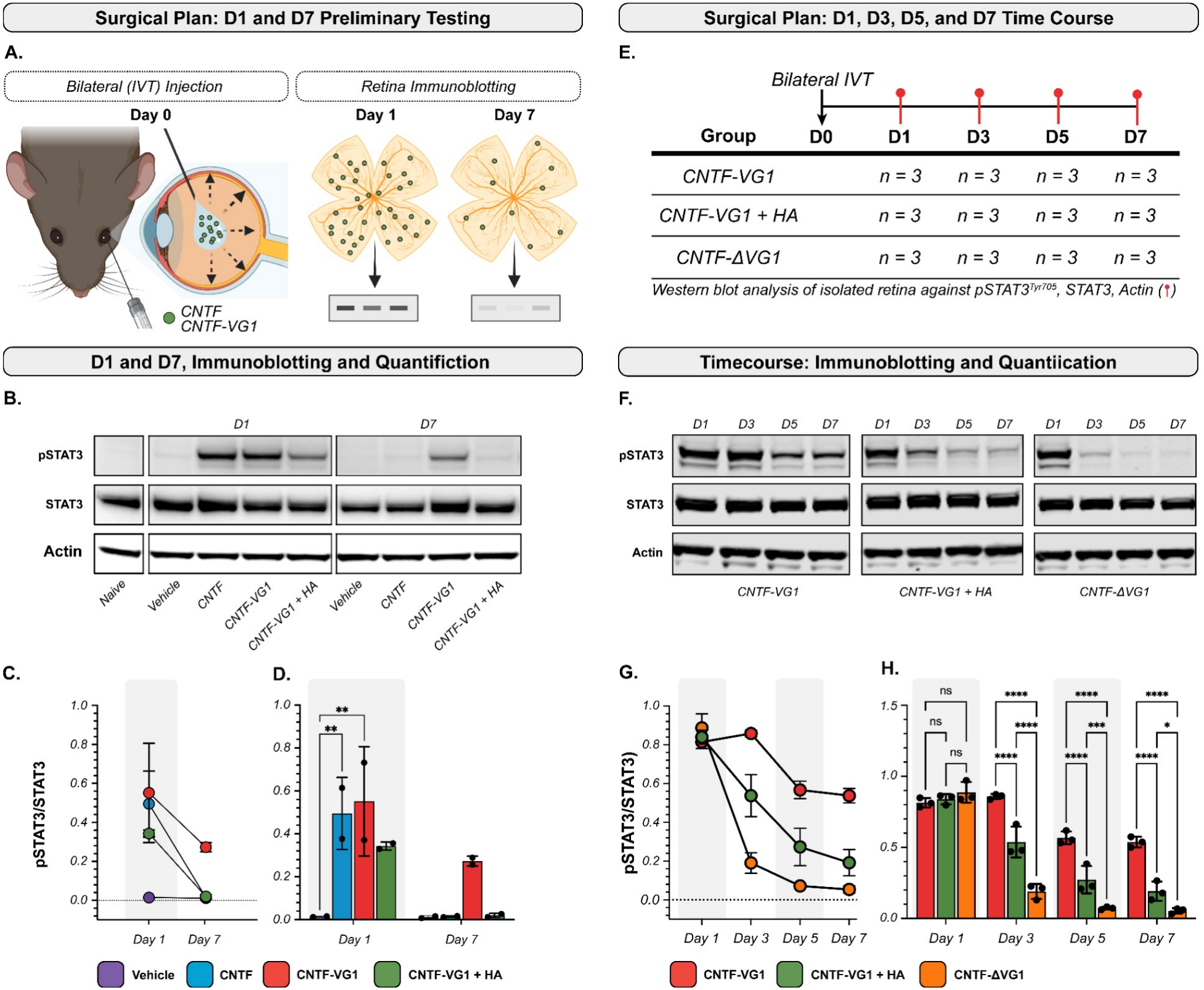
*Pharmacodynamic study of CNTF, CNTF-VG1, and CNTF-VG1 + HA in un-injured mice*. (A) Schematic presentation of experimental plan; un-injured mice eyes are injected either with vehicle (PBS; negative control), CNTF, CNTF-VG1 or CNTF-VG1 + HA, sacrificed, and homogenized retina are measured for pSTAT3^Tyr705^, STAT3, and Actin with immunoblotting at 1 and 7 days post-IVT injection. (B) Representative immunoblotting at Days 1 and 7. (C) Line graph and (D) tabulated pSTAT3^Tyr705^/STAT3 (normalized to actin). (E) Follow-up time course study summary of time points (Day 1, 3, 5, and 7), groups (CNTF-VG1, CNTF-VG1 + HA, and CNTF-ΔVG1), and animal breakdown. (F) Representative immunoblotting. (G) Line graph and (H) tabulated pSTAT3^Tyr705^/STAT3 (normalized to actin). Statistics: Two-way ANOVA, mixed model, multiple comparisons with post-hoc Tukey correction; data presented as mean ± SD; α = 0.05; * p < 0.05; ** p < 0.01; *** p < 0.001; **** p < 0.0001; ns = not significant; n = 3 (•).

CNTF-VG1 at Day 1, compared to naive and vehicle controls, suggests that our molecules are penetrating the inner limiting membrane (ILM) and reaching cells within the retina. Our results suggested that CNTF-VG1 has a higher JAK/STAT3 activation at D7 post-injection (0.272 ± 0.02) compared to CNTF (0.014 ± 0.004) and CNTF-VG1 + HA (0.019 ± 0.009), suggesting CNTF-VG1 may have a longer ocular retention window than our other groups. This goes against our expectation, as we originally hypothesized that co-formulation would extend the ocular retention time due to the combined effects of increased binding partners and greater protein stability conferred by co-formulation.

To probe this result and better understand the kinetics of our constructs, we conducted a follow-up study with a few procedural changes. First, we opted to investigate more granular timepoints–D1, D3, D5, and D7–to better understand the relative timelines of JAK/STAT3 activation within our different groups. Furthermore, to account for any differences in kinetics of diffusion potentially arising from the difference in hydrodynamic radius (R_H_), we introduced an additional CNTF-VG1 construct with double point mutations (D197A, Y208A) in the VG1 domain to ablate HA binding: CNTF-VG1^ΔD197A,ΔY208A^ (CNTF-ΔVG1; Figure S10A-D). Our group has previously reported that the individual mutations, D197A and Y208A, can significantly reduce HA-binding affinity while also affording increased thermal stability.^46^ To maximize this effect we designed and expressed a variant with both mutations (CNTF-ΔVG1), which possesses a near exact M_w_ match and bioactivity to CNTF-VG1 without measurable binding to HA (Figure S10E,F). A summary table of these groups and newly selected time points for our follow-up study is available in Figure 4E.

Representative immunoblotting for CNTF-VG1, CNTF-VG1 + HA, and CNTF-ΔVG1 for D1, 3, 5, and 7 are presented in Figure 4F; a line plot and tabulated statistics quantifying pSTAT3^Tyr705^/STAT3 can be found in Figure 4E and Figure 4G, respectively. Our second PD study had two significant results. First, we can confirm that VG1, specifically, significantly increases the JAK/STAT3 activation window of CNTF in murine eyes as indicated the CNTF-VG1 group having significantly (****, p < 0.0001) higher JAK/STAT3 activation, at all tested time points, compared to the CNTF-ΔVG1 group. Secondly, our study confirms the previous observation that the activation window of JAK/STAT3 is not only significantly decreased in our HA co-formulation group at day 7 compared to CNTF-VG1, but that this significant difference begins as early as day 3, with the overall JAK/STAT activation profile (Figure 4G) landing between CNTF-VG1 and CNTF-ΔVG1 over the course of the study. In summary, these data provide clear evidence that VG1, through HA-binding, can be used to increase the ocular retention and downstream activation of JAK/STAT3 in mice retina.

### CNTF is neuroprotective against reactive oxygen species-(ROS) induced death in primary mouse cortical neurons in an acute H_**2**_**O**_**2**_ cytotoxicity model

There is a growing body of evidence that ROS may be a critical modulator of RGC loss in glaucoma.^48^ When exposed to ROS, the accumulation of cell body and axonal damage initiates apoptosis through activation of cleaved caspases,^49^ which has been identified as one of the primary pathways of RGC loss in experimental glaucoma.^22,50^ CNTF has been proposed as a potential means to protect RGC’s through restoration of anti-oxidant levels^51^ and expression of anti-apoptotic factors in the retina.^52^ To test this *in vitro*, we quantified cell death in CNTF pre-treated primary mouse cortical neurons following exposure to chronic (continuous) or acute (short-term, <1 hr.) H_2_O_2_.

First, we established a relevant H_2_O_2_ treatment range by monitoring cell death over 72 hours through live-cell imaging using a nuclear-dye in MCN cultures treated with H_2_O_2_ (0 - 250 μM). Our results show that cytotoxicity (baseline: 20 ∼ 30% cell death for 0 μM H_2_O_2_ untreated condition) can be tuned over the range of approximately 60 ∼ 100% when exposed to 10 ∼ 100 μM H_2_O_2_ in a dose-dependent manner (Figure S11A,B; representative images: Figure S11C). Interestingly, we only could achieve a baseline ∼60% cell death, as below [H_2_O_2_] ≈ 10 μM (e.g., 1, 5 μM H_2_O_2_) we did not observe a significant increase in cell cytotoxicity.

To evaluate CNTF neuroprotection, we pre-treated MCN cultures for ∼24 hours with 100 ng mL^-1^ CNTF and measured cell death over the course of 3 days of chronic (continuous) 50, 25, and 10 μM H_2_O_2_ exposure. A schematic of our testing schema can be found in Figure S11D. However, in the chronic exposure condition we found no evidence of neuroprotection at any of the tested conditions (Figure S11E,F). We hypothesized that this lack of efficacy may be due to excessively high cytotoxic pressure, with cytotoxicity > 60% being beyond CNTF’s protective potential. To further probe this possibility, we developed an acute model by leveraging catalase as an ‘off switch,’ to harmlessly convert H_2_O_2_ into water and oxygen following timed H_2_O_2_ exposures.

We first validated that catalase could protect MCN’s against H_2_O_2_-induced and found that pre-treating MCN cultures with catalase completely ameliorates any cytotoxicity over our complete range of 0 - 250 μM H_2_O_2_ (Figure S12). Extending this approach, we tested timed exposures to H_2_O_2_ (15, 30, 45, and 60 min) to both 100 and 50 μM H_2_O_2_ (Figure S13A, B) alongside chronic (no catalase), untreated (no H_2_O_2_) and catalase pre-treated (1 hr. catalase pre-treated) control groups. Our findings show that by leveraging this acute exposure we gain much more granular control over viability. The 100 μM H_2_O_2_ condition has an extended tunable range from ∼40% to 100% cell death in ∼15% intervals, while the 50 μM H_2_O_2_ condition can be tuned similarly, but up to 80% cell death. Given the wider range of tuneability in the 100 μM H_2_O_2_ exposure condition, we adapted our chronic treatment strategy and measured cell death following acute exposures to 100 μM H_2_O_2_ in cultures with and without CNTF pre-treatment (treatment plan schematic: Figure 5A) and found that in the 30-minute condition, specifically, there is a significant (*, p < 0.05), sustained, decrease in cytotoxicity at most timepoints in cells pre-treated with CNTF compared to non CNTF-treated cells (Figure 5B, C; Summary of all acute 15-, 30-, 45-, and 60-minute and chronic conditions are provided in Figure S14). Importantly, while this finding just hits the threshold for significance when factoring in multiple runs, the trends were repeatable with varying significance across all our individual runs (Figure S15) suggesting its repeatability. Taken together, our data suggests that CNTF may be neuroprotective against ROS-induced neuronal death.

**Figure 5.**
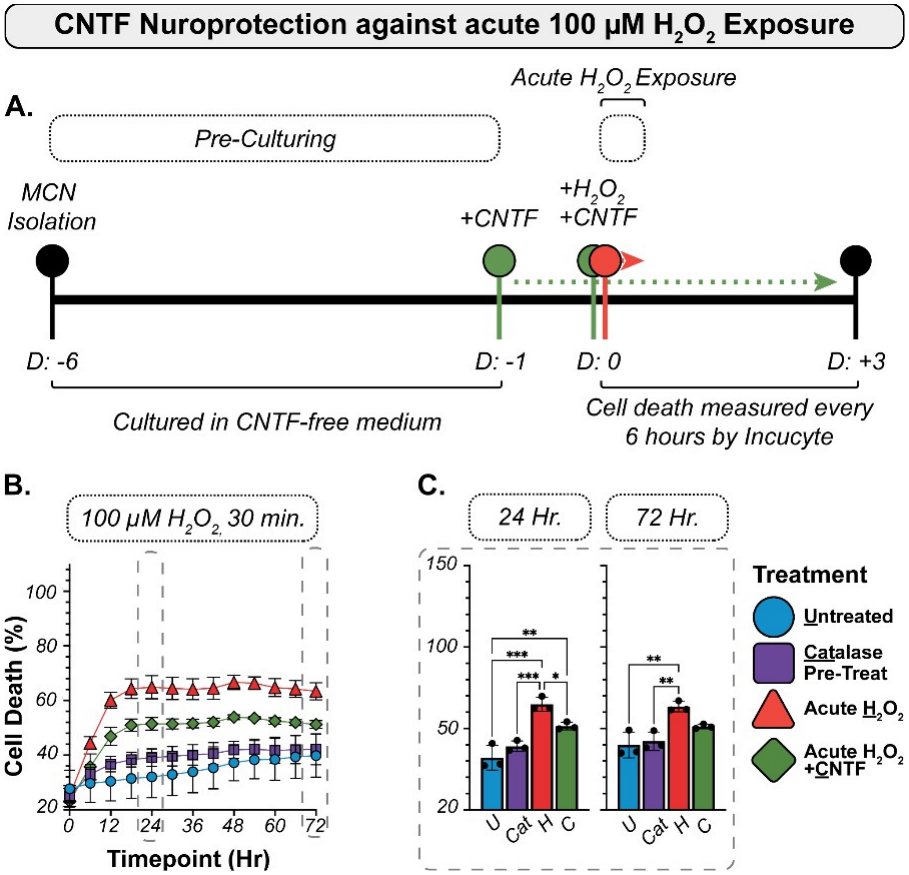
*Primary mouse cortical neuron H*_*2*_*O*_*2*_ *neuroprotection assay*. (A) Schematic representation of acute H_2_O_2_ neuroprotection assay. (B) Average time course of mouse cortical neuron (MCN) cell death following 30 min exposure to 100 μM without CNTF (Acute H_2_O_2_) and with CNTF pre-treatment (Acute H_2_O_2_ + CNTF) alongside untreated and catalase pre-treated controls Statistics on the time course graphs indicate significant differences between Acute H_2_O_2_ and Acute H_2_O_2_ + CNTF conditions only. (C) Summary statistics for both 24- and 72-hour time points are provided, for reference with statistics for all groups indicated. Statistics: Mixed model Two-Way ANOVA, multiple comparisons, with post-hoc Tukey correction test; data presented as mean ± SD; α = 0.05; * p < 0.05; ** p < 0.01; only significant differences are shown. n = 3 (•).

## DISCUSSION

Neuroprotection, in addition to reducing IOP, may significantly improve patient outcomes in glaucoma. In this framework, instead of addressing individual assaults (*e*.*g*., inflammation, ischemia, increased ROS, and elevated IOP) therapies are designed to make RGCs more resistant to cell death. However, because the eye has numerous physiological barriers to drug penetration,^53^ LAD strategies that provide consistent long-term bioavailability in the retina remain elusive. To address this limitation, we explored HA-binding *via* direct fusion with VG1 as a potential LAD strategy to extend the therapeutic window of potent neurotrophic factor CNTF following IVT injection. Our hypothesis is that a CNTF-VG1 fusion would have increased ocular retention through non-covalent interactions with endogenous HA in the vitreous, extending the residence time of CNTF, and therefore its window of potential neuroprotection (Figure 1). Our own group leveraged a similar strategy for augmenting brain retention of antibodies,^40^ however to our knowledge this is the first manuscript investigating the use of VG1 as a potential LAD technology for extended delivery of neurotrophic factors to the eye.

Our *in vitro* results clearly show that VG1 imparts strong HA-binding (*K*_*D*_ = ∼12 nM; Figure 2D, Figure S2A,B) to CNTF that further translates to a ∼50X reduction in diffusivity when exposed to primary vitreous *ex vivo* compared to PBS controls (Figure S7). Importantly, we also provide evidence that this can be done without significantly impacting its receptor binding (Figure 2E, Figure S2C,D) or activation of the critical neuroprotective JAK/STAT3 pathway in hESC-RGC (Figure S3). From a feasibility standpoint, our work goes on to show strong evidence that this is the case: both of our HA-binding groups, CNTF-VG1 and CNTF-VG1 + HA co-formulation, show significant increases in the window of neuroprotective JAK/STAT3 activation compared to CNTF and the non-binding CNTF-ΔVG1 mutant over all time points tested in uninjured mice eyes (Figure 4). To further validate this molecule as a potential therapeutic agent of interest we further developed our own variant of an H_2_O_2_ neurotoxicity assay that showed in short-term exposures CNTF offers, albeit modest, neuroprotection against ROS-induced neuronal death (Figure 5). A previous study found a similar result using rat CNTF in primary rat RGC cultures, however given the known difference in potency between human and rat CNTF^54^ we aimed to verify this effect and its specific conditions with our molecule. As a whole, this study sets the groundwork for a strategy to further leverage CNTF’s neuroprotective potential in the fight against glaucoma.

However, our work also highlights a key consideration for ECM-binding, broadly, as a strategy for ocular delivery: HA-binding, and likely any ECM-binding moiety, has the potential to induce unfavorable protein-protein interactions and induce aggregation. We showed very clearly that CNTF-VG1 dramatically precipitates in low-HA solutions, and that this is further exaggerated at physiological temperatures (Figure 3). While we do not have conclusive evidence, we hypothesize that this is due to the hydrophobic CNTF molecules being forced into a pseudo-high concentration as a side-effect of competitive binding for a limited number of free-HA molecules. The fact that observable precipitation is almost entirely eliminated in the presence of excess-HA, across both room temperature and physiological temperatures (Figure 3D and Figure 3H, respectively), provides credence to this theory, as providing an excess of HA-binding partners may both prevent this pseudo-high concentration and affect the overall hydrophilicity of CNTF in its bound-state. From a translation perspective, HA delivery in the form of artificial vitreous is already being investigated pre-clinically and has evidence of being both biocompatible and non-disruptive to optical properties of the vitreous, suggesting that HA co-formulation may be a viable strategy in this application.^55,56^ However, while these precipitation effects can be ameliorated by co-formulation with HA, this approach has the potential to significantly alter the molecules pharmacokinetics as we observed in our own study where the HA co-formulation group had a JAK/STAT3 activation profile that landed almost precisely between CNTF-VG1 and the non-binding CNTF-ΔVG1 groups (Figure 4E-H).

Presently, we do not have a complete understanding of the mechanism behind this attenuation. However, we speculate that it may be because in the HA co-form group CNTF-VG1 is paired with an excess of relatively (compared to endogenous) low M_w_ HA that exfiltrates it out of the eye faster than the CNTF-VG1-only group in which a soluble, non-aggregated, portion of CNTF-VG1 may still be able to bind with endogenous HA and enable increased retention. However, these findings also suggest another useful implication of HA pre-complexing that has not previously been explored: it may allow researchers to ‘tune’ the PK/PD window by mixing HA-binding molecules with HA prior to injection. This would need to be balanced with its effect on viscosity; larger Mw HA species may, in this proposed mechanism, slow diffusion but will have the tradeoff of increasingly making the injection solution increasingly viscous.

One could make the case that the challenge of protein precipitation and aggregation could be engineered out. Already others have shown that certain amino acid mutations in WT CNTF can, for example, significantly improve its stability and overall solubility.^57^ However, this is non-trivial and would require molecule-specific considerations as even single point mutations can have sweeping knock-on effects to other molecule properties. In the case of CNTF, for example, single point mutants can ablate receptor binding with one of its three receptors–gp130, LIFRβ, and CNTFRα–and significantly alter its bioactivity.^58,59^ Importantly, all of these effects would also need to be considered in the context of a VG1 fusion. In our own hands, we found that fusion with VG1 could qualitatively improve the overall solubility of CNTF. At our dosing concentrations (CNTF = 2 mg mL^-1^; CNTF-VG1 = 5.08 mg mL^-1^), we observed that CNTF could be pushed to as high as 2 mg mL^-1^ through concentration at 4 °C, but that it was metastable, with evidence of slow precipitation over the days that followed (Figure S16). Conversely, CNTF-VG1 remained fully soluble with no visual evidence of precipitation up to concentrations > 15 mg mL^-1^ when stored at 4 °C (data not shown).

Furthermore, while in-depth stability characterization is beyond the scope of our current study, we also observed that CNTF-VG1 is susceptible to significant long-term aggregation in storage conditions, with as much as a 35% HMWS forming when stored as a liquid formulation after a year at 4 °C (Figure S17A-D). This is in stark contrast to VG1 where we have seen high stability (>95% monomeric species) even after 3 years in storage under identical conditions (Figure S17E). Given the identified aggregation and solubility challenges highlighted previously with our CNTF-VG1 molecule, we suspect this may be tied to the pendant CNTF. These data and observations are practical realities that must be confronted when considering neurotrophic factor delivery generally but are heightened when the delivery mechanism artificially increases the propensity of protein-protein interactions as with HA-binding. In short, the viability of ECM-binding as a strategy is molecule specific, but our work presents a framework to evaluate potential molecules of interest early in development.

Our study has numerous open questions which were not adequately addressed here, and we outline some of them presently here. First, our *in vivo* evaluation methods suggest an extended JAK/STAT3 activation window, but it does not provide insight into where our molecule is sequestering, or the precise mechanism it is acting under to achieve that extended window. Based on the readout from our CNTF-ΔVG1 data we know HA-binding is pivotal in increasing ocular retention in this model. However, given the low [HA] in mice vitreous suggested by our vitreous HA ELISA data (Figure S4), only a small fraction may actually be in the vitreous while the rest is distributed to other HA-rich tissues in the eye: ocular blood vessels, aqueous humor, cornea, retina, basal layer of the inner limiting membrane, or others.^60–62^ Latent in our data is another consideration for ocular researchers: mice, rats, and even rabbits, may not be ideal candidates for studying the effect of HA-binding in the eye as these smaller animal models have significantly lower vitreous [HA] than their larger-mammal counterparts–pigs, monkeys, and humans^63^–rendering cross-species comparisons challenging, at best.

An additional unanswered question is whether CNTF-VG1 aggregates, as we predicted it would, post-injection and how that factors into the final JAK/STAT3 window we observed in relation to the HA co-formulation group. As alluded to earlier, one possible explanation supported by our FCS data (Figure S7) is that a soluble fraction of CNTF-VG1 was able to bind to the relatively high M_w_ endogenous HA which would have, therefore, a slower diffusion than the molecules in our HA co-formulation group where there is an excess of more mobile, lower M_w_ HA species. However, without a means to carefully track our molecules’ physical location post-injection, this cannot be easily addressed. Beyond this, still, other questions as to the retinal biodistribution and confirmation as to which cells are actively getting exposed to CNTF in our study remain open. Numerous cells in the retina beyond RGCs have adequate receptors to engage with CNTF, such as photoreceptors, Müller glial cells, and retinal pigment epithelial cells, and the relative contribution to our readout is unclear as our methodology analyzed the retina in whole.

Taken together, this paper provides a compelling case that HA-binding may be a potential strategy for gaining extended ocular retention of neurotrophic factors following IVT. While our work was focused on ocular delivery for applications in glaucoma, there are numerous other HA-rich organs–skin, cardiovascular tissue, blood vessels, the brain and spinal cord, and others–that may also benefit from the extended retention provided by ECM-binding moieties. While there are potential challenges in and around aggregation stemming from that method of action, careful target and molecule selection may make the most of this unique LAD opportunity.

## CONCLUSION

Direct fusion of neuroprotective ciliary neurotrophic factor (CNTF) with HA-binding G1 domain of versican (VG1; CNTF-VG1) imparts high affinity HA-binding (*K*_*D*_: ∼10 nM) without significantly altering CNTF’s native bioactivity or down-stream JAK/STAT3 signaling pathway. This translates to a near ∼50X decrease in the diffusivity of CNTF-VG1 in samples of primary vitreous compared to PBS. We show that HA-binding can induce interactions that lead to significant aggregation in cases where [CNTF-VG1] >> [HA], however we go on to show that this aggregation can be ablated through co-formulation with excessive HA in addition to conferring increased thermal stability. Additionally, we show that *ex vivo* this co-formulation approach significantly reduces precipitation in primary pig and rat vitreous. Through a two-part *in vivo* study, we show that: 1) VG1 significantly increases the activation window of JAK/STAT3 compared to non-binding controls and that, interestingly, co-formulation with HA attenuates the window of JAK/STAT3 activation. We close our work with an *in vitro* H_2_O_2_ cytotoxicity assay in primary mouse cortical neurons and show that CNTF treated cells are conferred with modest resistance to ROS-induced neurotoxicity.

## ASSOCIATED CONTENT

*The associated supporting information is summarized below:*

**Figure S1**: *CNTF, CNTF-VG1, and VG1 Schematics*.

**Figure S2**: *Representative SPR and TF-1 proliferation curves with tabulated summary data*

**Figure S3**: *JAK/STAT3 activation in hESC-RGC and primary mouse cortical neuron (MCN) cell cross-reactivity testing*

**Figure S4**: *Vitreous HA ELISA assay*

**Figure S5**: *CNTF-VG1 and HA M*_*w*_ *and concentration stability screen (23* ^*o*^*C)*

**Figure S6**: *CNTF-VG1 and HA M*_*w*_ *and concentration stability screen (37* ^*o*^*C)*

**Figure S7**: *CNTF-VG1 diffusivity measurement in fluorescence correlation spectroscopy*

**Figure S8**: *HA co-formulation effect on vitreous stability of CNTF-VG1 testing protocol*

**Figure S9**: *Background membrane imaging (BMI) particle masking example images*

**Figure S10**: *Attenuated HA-binding CNTF-ΔVG1 mutant HA-binding kinetics, bioactivity*

**Figure S11**: *CNTF neuroprotection against chronic H*_*2*_*O*_*2*_ *exposure in MCN culture*

**Figure S12**: *Effect of catalase pre-treatment on MCN exposed to H*_*2*_*O*_*2*_.

**Figure S13**: *Acute H*_*2*_*O*_*2*_ *treatment viability curves for 100, 50 µM H*_*2*_*O*_*2*_ *treatment conditions*.

**Figure S14**: *CNTF neuroprotection against acute (15, 30, 45, and 60 min*.*) and chronic 100 µM H*_*2*_*O*_*2*_ *exposure in primary mouse cortical neuron cultures (n = 3)*

**Figure S15**: *Individual biological replicates for CNTF neuroprotection assay against acute 100 µM H*_*2*_*O*_*2*_ *exposure in MCN*

**Figure S16**: *CNTF and CNTF-VG1 turbidity images for at-dose concentrations (4* ^*o*^*C)*

**Figure S17**: *CNTF and CNTF-VG1 aggregation over time under storage conditions*

## AUTHOR INFORMATION

**Riley A. Suhar** - *Pharmaceutical Development, Drug Delivery, Genentech, 1 DNA Way, South San Francisco, California 94080, United States orcid*.

**Xiaowu Gu** - *Research and Early Development, Neuroscience, Genentech, 1 DNA Way, South San Francisco, California 94080, United States orcid*.

**Alexandra Robles** - Purification, Microbiology and Virology, *Genentech*, 1 DNA Way, South San Francisco, California 94080, United States

**Robert F. Kelley** - *Pharmaceutical Development, Drug Delivery, Genentech, 1 DNA Way, South San Francisco, California 94080, United States orcid*.

**Devin B. Tesar** - *Pharmaceutical Development, Drug Delivery, Genentech, 1 DNA Way, South San Francisco, California 94080, United States*

## ACKNOWLEDGEMENTS

R.A. Suhar acknowledges M. Jeanne, P. Teotia, and F. Abbas for their thoughtful discussion, patience, and mentorship in the early stages of this work; RA. Suhar further would like to extend a special thank you and gratitude to C. Hoogenraad for his immense support during difficult project transitions–he went above and beyond what most would. Thank you. It is not forgotten. R.A. Suhar further acknowledges the long-term helpful discussion and feedback from the members of the PTD Drug Delivery Group, Neuroscience, and the assorted personnel across the SSF campus who have intersected with this body of work–indirectly or otherwise–for their curiosity-driven feedback and helpful push when the work was stuck, it would not have been possible without all of them. The authors extend their gratitude to the reviewers, internal and external, who gave their time to make this work the best version it can be in the name of scientific rigor. Lastly, we acknowledge the Genentech Postdoctoral Program for the financial support and non-material contributions that provided the opportunity for this work to be completed.

## Notes

### Competing Interest Statement

The authors have declared no competing interest.

