## SUPPLEMENTAL INFORMATION for "Extending intravitreal retention of CNTF via fusion to the HA-binding protein VG1"

**SUMMARY:**

**Figure S1**: *CNTF, CNTF-VG1, and VG1 schematics.*
**Figure S2**: *Representative SPR and TF-1 proliferation curves with tabulated summary data.*
**Figure S3**: *JAK/STAT3 activation in hESC-RGC and primary mouse cortical neuron (MCN) cell cross-reactivity testing*
**Figure S4**: *Vitreous HA ELISA assay*
**Figure S5**: *CNTF-VG1 and HA $M_w$ and concentration stability screen (23 ºC).*
**Figure S6**: *CNTF-VG1 and HA $M_w$ and concentration stability screen (37 ºC).*
**Figure S7**: *CNTF-VG1 diffusivity measurement in fluorescence correlation spectroscopy.*
**Figure S8**: *HA Co-formulation effect on vitreous stability of CNTF-VG1 testing protocol.*
**Figure S9**: *Background membrane imaging (BMI) particle masking example images.*
**Figure S10**: *Attenuated HA-binding CNTF-ΔVG1 mutant HA-binding kinetics, bioactivity.*
**Figure S11**: *CNTF neuroprotection against chronic $H_2O_2$ exposure in MCN culture.*
**Figure S12**: *Effect of catalase pre-treatment on MCN exposed to $H_2O_2$*
**Figure S13**: *Acute $H_2O_2$ treatment viability curves for 100, 50 µM $H_2O_2$ treatment conditions.*
**Figure S14**: *CNTF neuroprotection against acute (15, 30, 45, and 60 min.) and chronic 100 µM H2O2 exposure in primary mouse cortical neuron cultures (n = 3) .*
**Figure S15**: *Individual biological replicates for CNTF neuroprotection assay against acute 100 µM $H_2O_2$ exposure in MCN.*
**Figure S16**: *CNTF and CNTF-VG1 turbidity images for at-dose concentrations (4 ºC).*
**Figure S17**: *CNTF and CNTF-VG1 aggregation over time under storage conditions.*

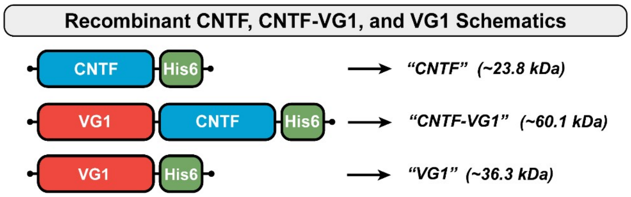


**Figure S1**: *CNTF, CNTF-VG1, and VG1 schematics.*
*Schematic representations of CNTF-his6 (CNTF), VG1-CNTF-his6 (CNTF-VG1), and VG1-his6 (VG1).*

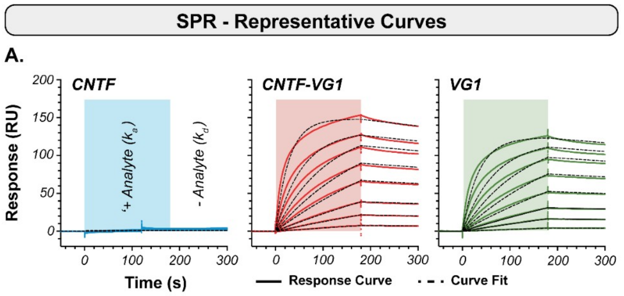


SPR Average Kinetics

B.

| Protein | $k_a$ ($M^{-1}$ $s^{-1}$) x $10^{-4}$ | $k_d$ ($s^{-1}$) x $10^5$ | $K_D$ (nM) |
|---|---|---|---|
| *CNTF* | - | - | - |
| *CNTF-VG1* | *16.2 ± 8.7* | *143.7 ± 60.4* | *11.7 ± 7.6* |
| *VG1* | *11.5 ± 2.6* | *450.8 ± 27.3* | *40.2 ± 8.0* |

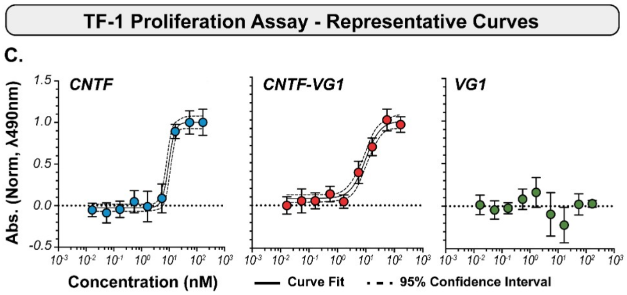


TF-1 Relative Bioactivity - Summary Statistics

D.

| Protein | $ED_{50}$ (nM) | $ED_{50}$ (ng $mL^{-1}$) |
|---|---|---|
| *CNTF* | *8.64 ± 0.75* | *206.6 ± 17.89* |
| *CNTF-VG1* | *11.53 ± 2.31* | *694.5 ± 139.03* |
| *VG1* | - | - |

**Figure S2**: *Representative SPR and TF-1 proliferation curves with tabulated summary data.*
Representative surface plasmon resonance (SPR) curves, with fits, for (A) CNTF (negative control, no-response), CNTF-VG1 and VG1 (positive control) constructs against immobilized hyaluronic acid (HA). (B) Tabulated results for the binding ($k_a$), dissociation ($k_d$) and equilibrium dissociation constant ($K_D$) kinetic factors; data shown as mean ± SD. (C) Representative TF-1 proliferation response curves for CNTF (positive control), CNTF-VG1, and VG1 (negative control, no response) with curve fit (Sigmoidal, 4PL). (H) Tabulated bioactivity results with interpolated $ED_{50}$ proliferation, presented as concentration (nM and ng $mL^{-1}$); data shown as mean ± SD.

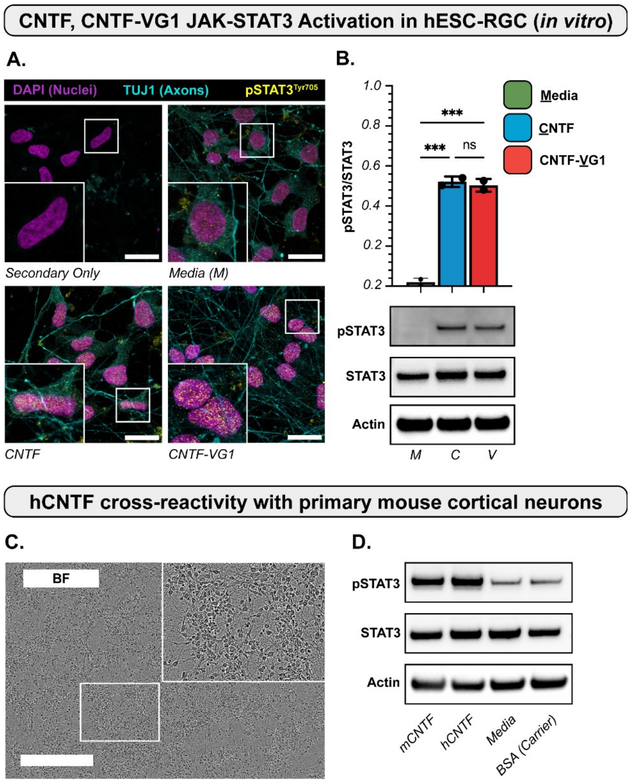


**Figure S3**: *JAK/STAT3 activation in hESC-RGC and primary mouse cortical neuron (MCN) cell cross-reactivity testing.* (A) Representative immunofluorescent images (DAPI (nuclei), TUJ1 (axons), and pSTAT3$^{Tyr705}$) of fixed human embryonic stem cell-derived retinal ganglion cells (hESC-RGC) treated for 30 minutes with media (negative control), CNTF, and CNTF-VG1 alongside a secondary-only control for reference (scale bar = 25 µm). (B) Tabulated pSTAT3$^{Tyr705}$/STAT3 from immunoblotting of treated hESC-RGC. (C) Representative bright field images of primary mouse cortical neurons (MCN) and (D) immunoblotting of MCN cultures treated with: mCNTF, hCNTF, media and 0.1% bovine serum albumin (BSA) carrier. Statistics: One-way ANOVA, multiple comparisons with post-hoc Tukey correction; data presented as mean ± SD; $\alpha = 0.05$; *** $p < 0.001$; ns = not significant; n = 2 (•).

**Vitreous HA Concentration - ELISA Assay**

**A.**

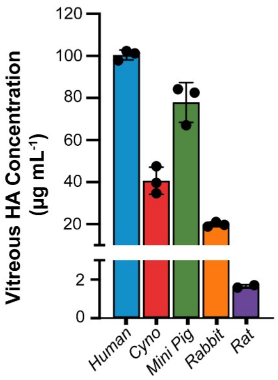


**B.**

| Species | Vitreous [HA] (µg mL$^{-1}$) |
|---|---|
| *Human* | *100.4 ± 3.1* |
| *Cyno* | *40.6 ± 6.4* |
| *Pig* | *77.9 ± 9.5* |
| *Rabbit* | *19.9 ± 1.1* |
| *Rat* | *1.6 ± 0.1* |

**Figure S4**: *Vitreous HA ELISA assay.*
(A) ELISA assay measuring hyaluronic acid content in human, mini pig (pig), cynomolgus macaque (cyno), rabbit, and rat vitreous (statistics not shown). (B) Tabulated HA ELISA results for reference.

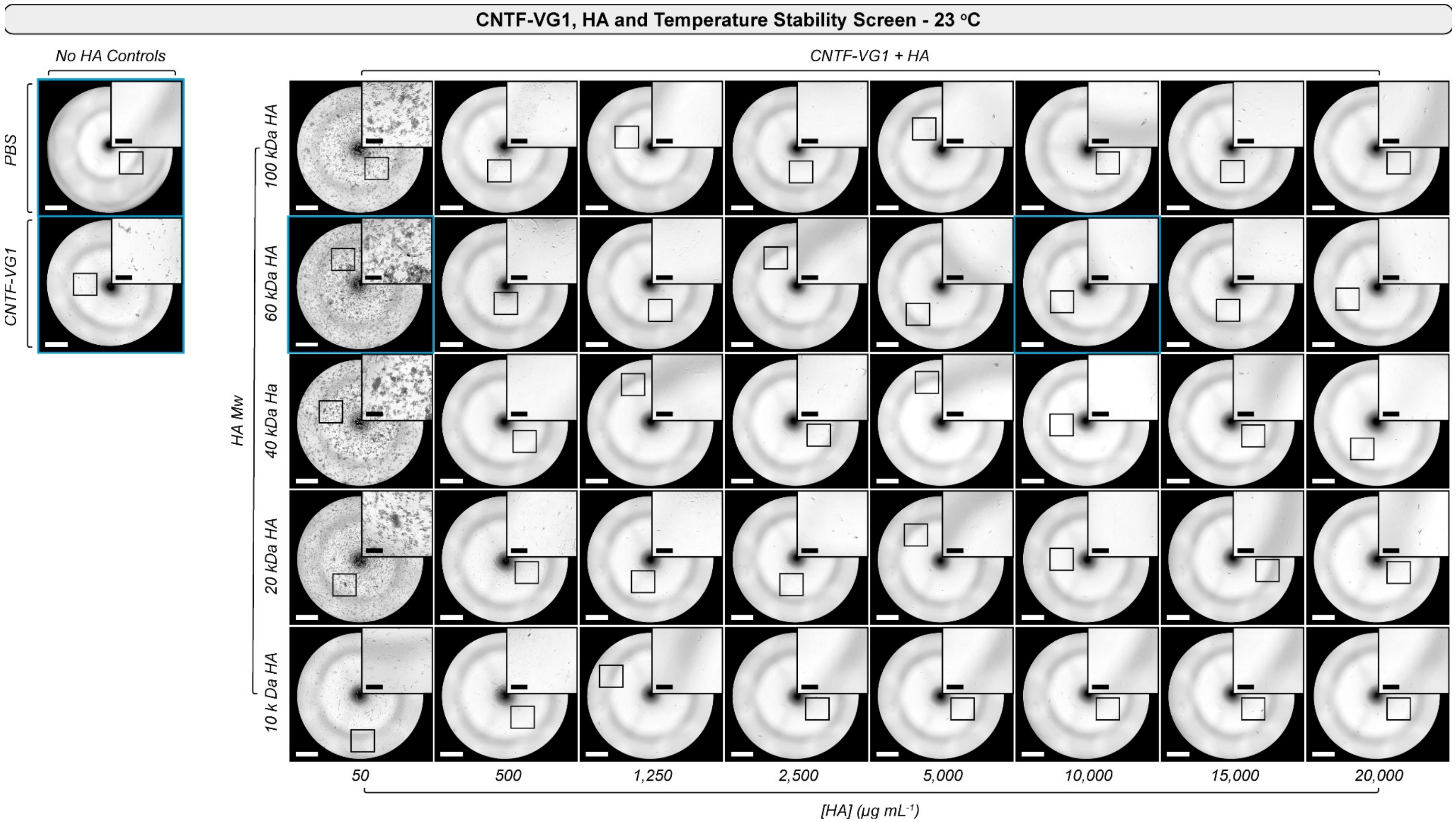


**Figure S5:** *CNTF-VG1 and HA $M_w$ and concentration stability screen (23 °C).*
Representative brightfield images from solutions of CNTF-VG1 (5.0 mg $mL^{-1}$, final) and a range of $M_w$ and [HA] (full HA $M_w$ range: 10 - 100 kDa; full [HA] range: 0 - 20,000 µg $mL^{-1}$) incubated for 24 hours at room temperature (23 °C) along with PBS and CNTF-VG1 only controls (white scale bar = 2-mm; black scale bar = 500 µm). The specific images that were reproduced in Figure 3A-D are outlined in blue.

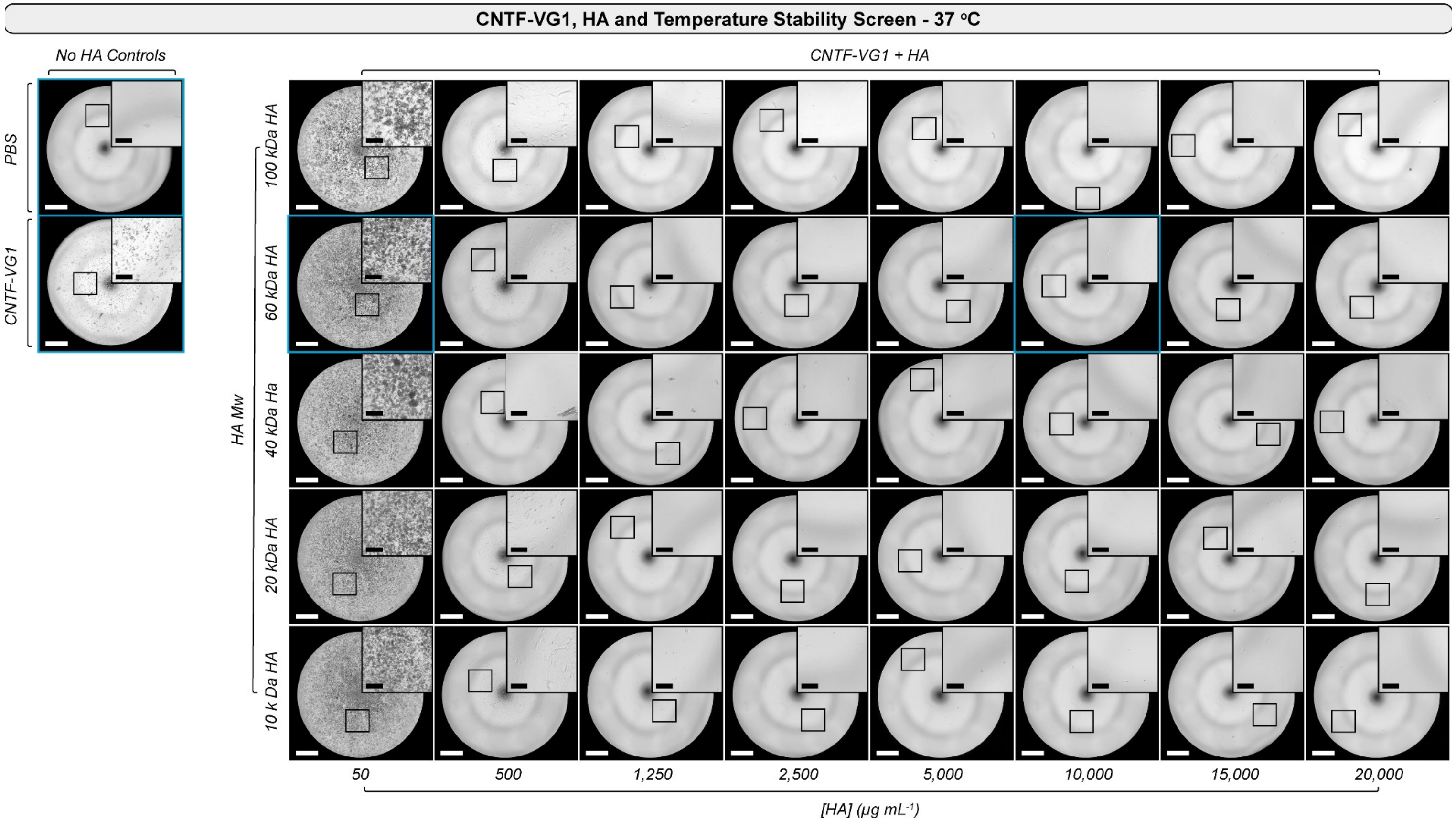


**Figure S6**: *CNTF-VG1 and HA $M_w$ and concentration stability screen (37 °C).*
Representative brightfield images from solutions of CNTF-VG1 (5.0 mg $mL^{-1}$, final) and a range of $M_w$ and [HA] (full HA $M_w$ range: 10 - 100 kDa; full [HA] range: 0 - 20,000 µg $mL^{-1}$) incubated for 24 hours at (37 °C) along with PBS and CNTF-VG1 only controls (white scale bar = 2-mm; black scale bar = 500 µm). The specific images that were reproduced in Figure 3E-H are outlined in blue.

**CNTF-VG1 Diffusivity in FCS**

**A.**

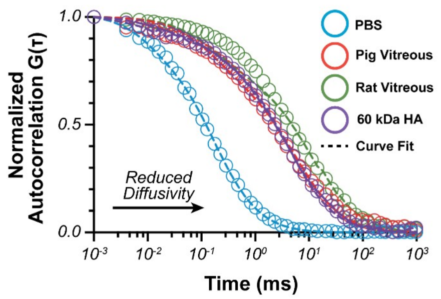


**B.**

| **Medium** | **D** (μm² s⁻¹) |
|---|---|
| *PBS* | *54.25* |
| *Pig Vitreous* | *1.62* |
| *Rat Vitreous* | *0.904* |
| *60 kDa HA* | *2.35* |

**Figure S7**: *CNTF-VG1 diffusivity measurement in fluorescence correlation spectroscopy.*
(A) Representative, normalized, fluorescence correlation spectroscopy (FCS) traces of CNTF-VG1-AF488 in highly dilute solutions (CNTF-VG1 << HA) of: PBS, pig vitreous, rat vitreous, and 60 kDa HA with curve fits. (B) Tabulated diffusivity (D; $\mu m^2\ s^{-1}$) measurements for each condition.

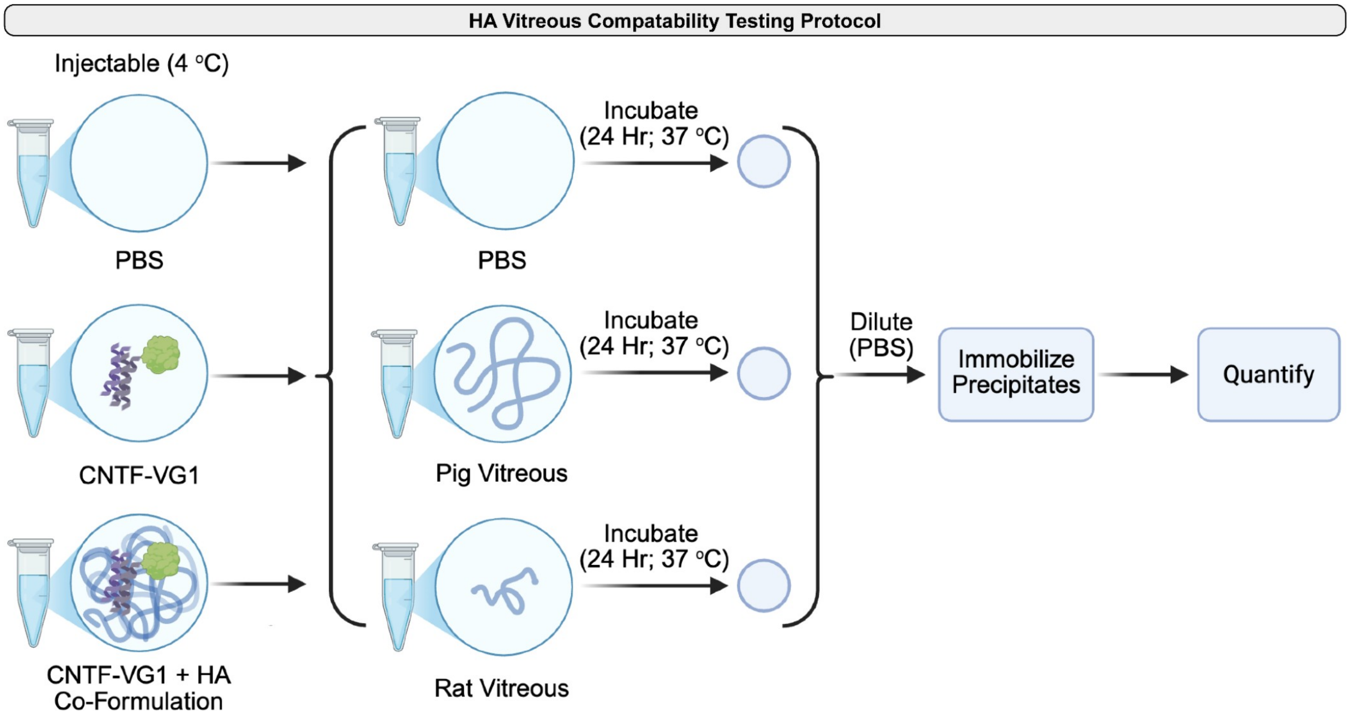


**Figure S8**: *HA co-formulation effect on vitreous stability of CNTF-VG1 testing protocol.*
Schematic outline of proposed vitreous compatibility assay. In short, solutions of PBS, CNTF-VG1 (5.0 mg mL$^{-1}$, final) and CNTF-VG1 + HA co-formulation (5.0 mg mL$^{-1}$ CNTF-VG1 final); 1% w/v 60 kDa HA, final) were pipetted (without mixing) into a pre-warmed (37 ºC) solutions of either PBS, pig vitreous or rat vitreous and incubated for 24 hours at 37 ºC. After 24 hours of incubation, samples were diluted (1:10, pre-warmed PBS) and precipitates were then immobilized and quantified using background membrane imaging. Created in *BioRender*. Suhar, R. (2026) https://BioRender.com/u0ycti6.

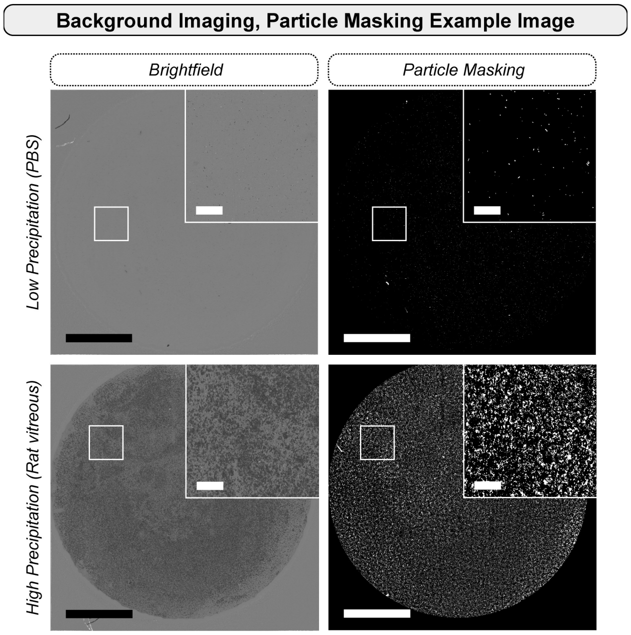


**Figure S9**: *Background membrane imaging (BMI) particle masking example images.*
Representative particle masking images for a low- (top row) and high- (bottom row) precipitation solutions (main scale bar = 2-mm, zoom-in scale bar = 500 µm).

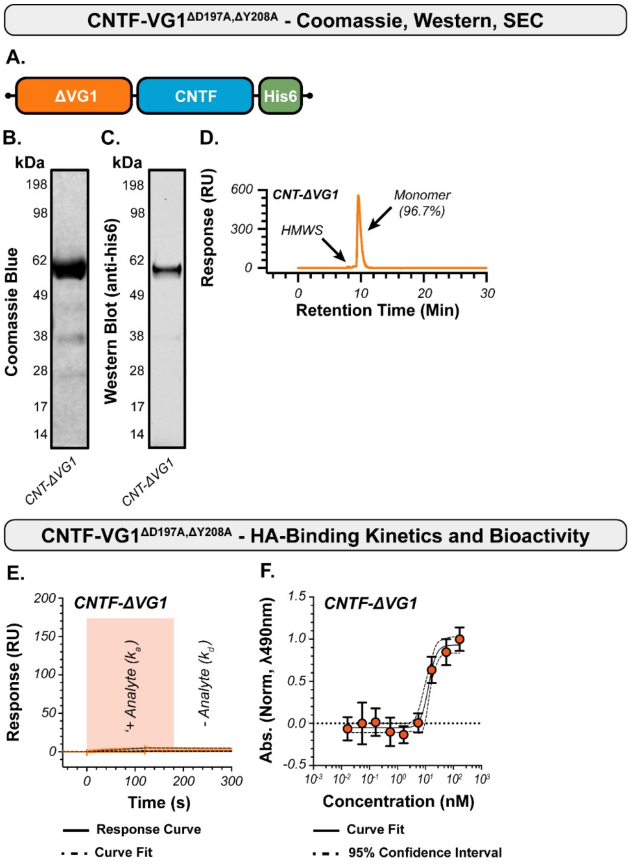


G.

| Protein | $K_D$ (nM) | $ED_{50}$ (ng mL$^{-1}$) |
|---|---|---|
| *CNTF-ΔVG1* | - | *10.04 ± 2.69* |

**Figure S10**: *Attenuated HA-binding CNTF-ΔVG1 mutant HA-binding kinetics, bioactivity.*
(A) Schematic representation of double point-mutant, non- HA-binding, CNTF-VG1$^{\Delta D197A,\Delta Y208A}$ (CNTF-ΔVG1). Representative (B) Coomassie stain (black and white), (C) western blot (anti-his6) and (D) size exclusion chromatography (SEC) trace for CNTF-ΔVG1. (E) Representative SPR curves, with fits, for CNTF-ΔVG1 against immobilized hyaluronic acid. (F) Representative TF-1 proliferation response curve for CNTF-ΔVG1 with curve fit (Sigmoidal, 4PL). (G) Summarized results for equilibrium dissociation constant ($K_D$; nM; no response) and $ED_{50}$ proliferation. Data shown as mean ± SD.

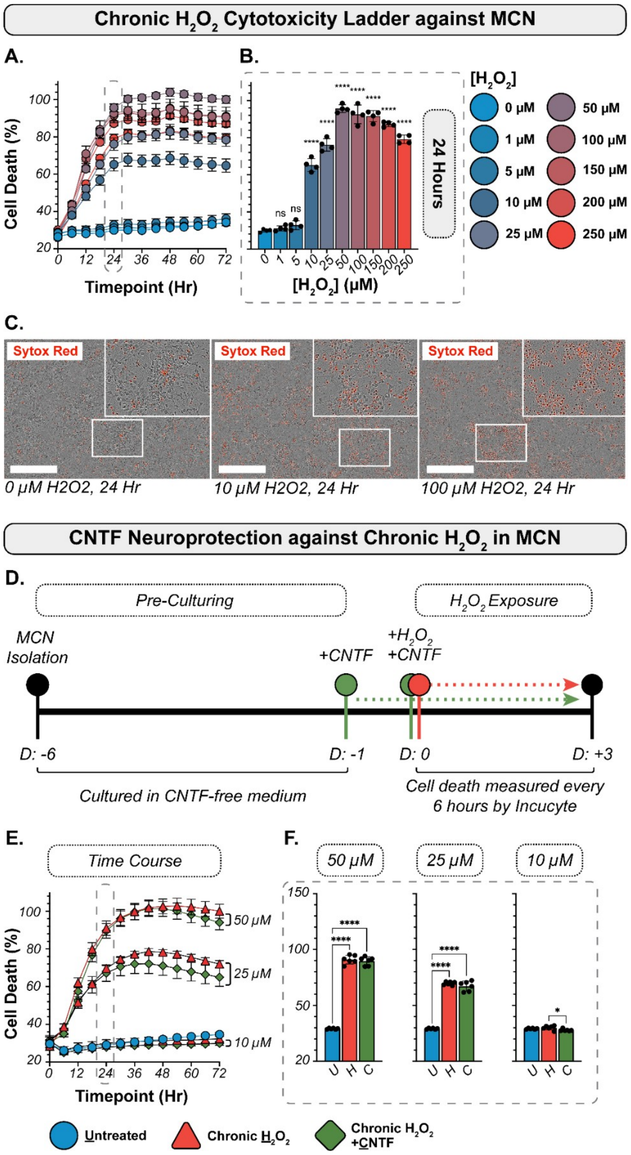


**Figure S11**: *CNTF neuroprotection against chronic $H_2O_2$ exposure in MCN culture.*
(A) Representative MCN viability curves (normalized to highest cell death) for a range of $H_2O_2$ (chronic; 0 - 250 μM) tested. (B) Viability at 24-hour time point with statistical comparisons references to the untreated (0 μM $H_2O_2$) condition. (C) Representative images (overlayed brightfield + Sytox Red) for 0, 10, and 100 μM $H_2O_2$ at 24-hour time-point (scale bar = 400 μm). (D) Chronic $H_2O_2$ neuroprotection assay outline schematic. (E) Time course readout of cell viability for 50, 25, and 10 μM $H_2O_2$ chronic exposure conditions with and without CNTF pre-treatment, plotted against untreated conditions. (F) Statistical comparisons at 24-hour timepoint for chronic 50, 25 and 10 μM $H_2O_2$ conditions. Statistics: Mixed model Two-Way ANOVA, multiple comparisons with repeated measures, with post-hoc Tukey correction test; data presented as mean ± SD; α = 0.05; * $p < 0.05$; *** $p < 0.001$; **** $p < 0.0001$; n = 4-6 (•).

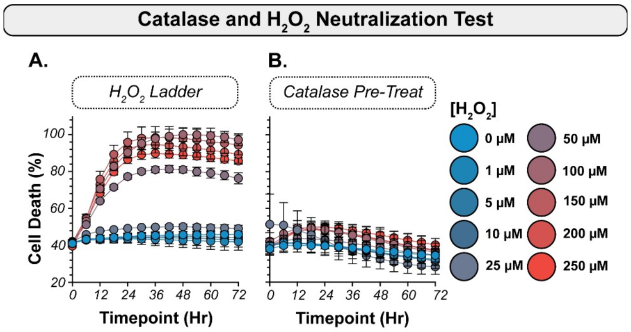


**Figure S12**: *Effect of catalase pre-treatment on MCN exposed to $H_2O_2$.*
(A) Representative MCN viability curves (normalized to highest cell death) for a range of [$H_2O_2$] (chronic; 0 - 250 µM) treatment conditions. (B) Representative MCN viability curves for a range of [$H_2O_2$] treatment conditions following pre-treatment (1 hr.) with catalase.

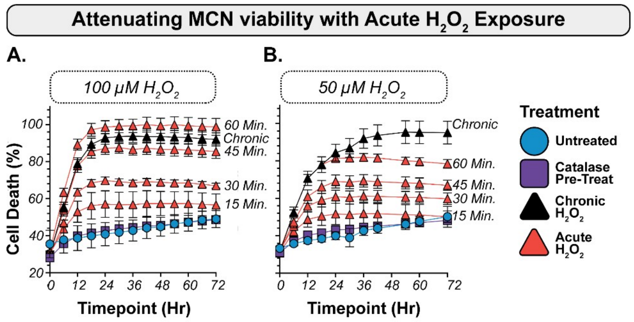


**Figure S13**: *Acute $H_2O_2$ treatment viability curves for 100, 50 µM $H_2O_2$ treatment conditions.*
Representative MCN viability curves (normalized to highest cell death for (A) 100 µM $H_2O_2$ and (B) 50 µM $H_2O_2$ conditions. For each condition, untreated (0 µM $H_2O_2$), catalase pre-treated (1 hour catalase pre-treatment followed by addition of $H_2O_2$) and 15-, 30-, 45-and 60-minute acute $H_2O_2$ exposure (timed exposure of $H_2O_2$ followed by addition of catalase) are shown.

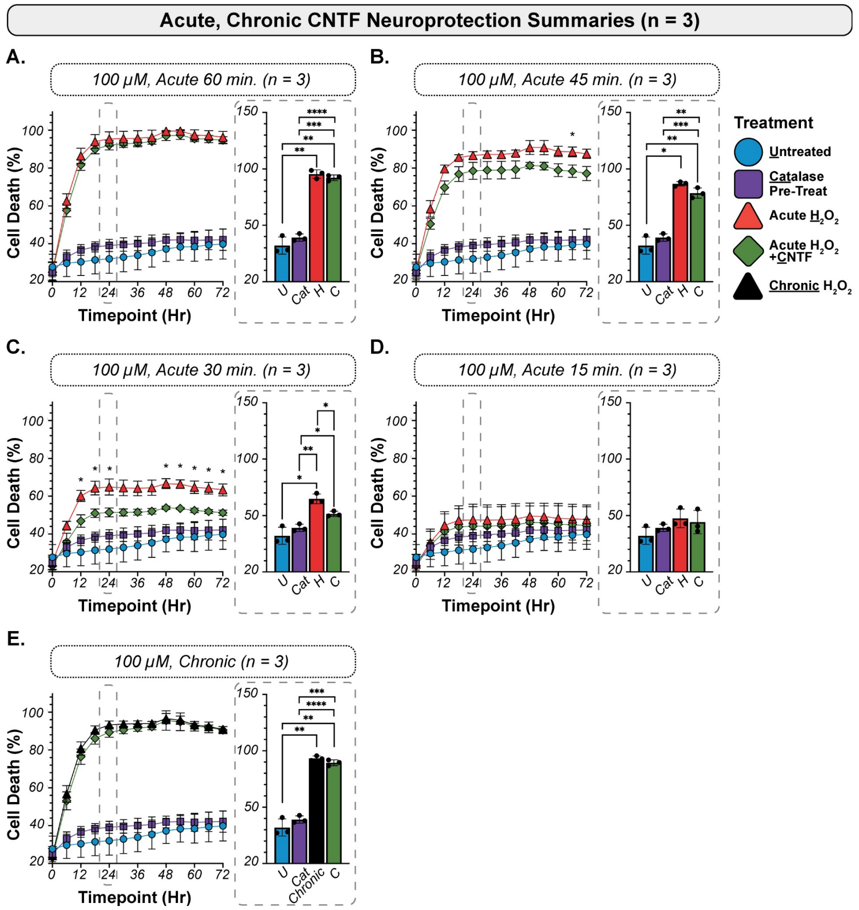


**Figure S14**: *CNTF neuroprotection against acute (15, 30, 45, and 60 min.) and chronic 100 µM $H_2O_2$ exposure in primary mouse cortical neuron cultures (n = 3).*
Summary (n = 3) of (A) 15 min., (B) 30 min., (C) 45 min., and (D) 60 min. acute and (E) chronic exposure conditions for 100 µM $H_2O_2$ in MCN cultures. Statistics on the time course graphs indicate significant differences between Acute $H_2O_2$ and Acute $H_2O_2$ + CNTF conditions only. Statistics: Mixed model Two-Way ANOVA, multiple comparisons with repeated measures, with post-hoc Tukey correction test; data presented as mean ± SD; α = 0.05; * $p < 0.05$; ** $p < 0.01$; *** $p < 0.001$; **** $p < 0.0001$; only significant differences are shown; n = 3 (•).

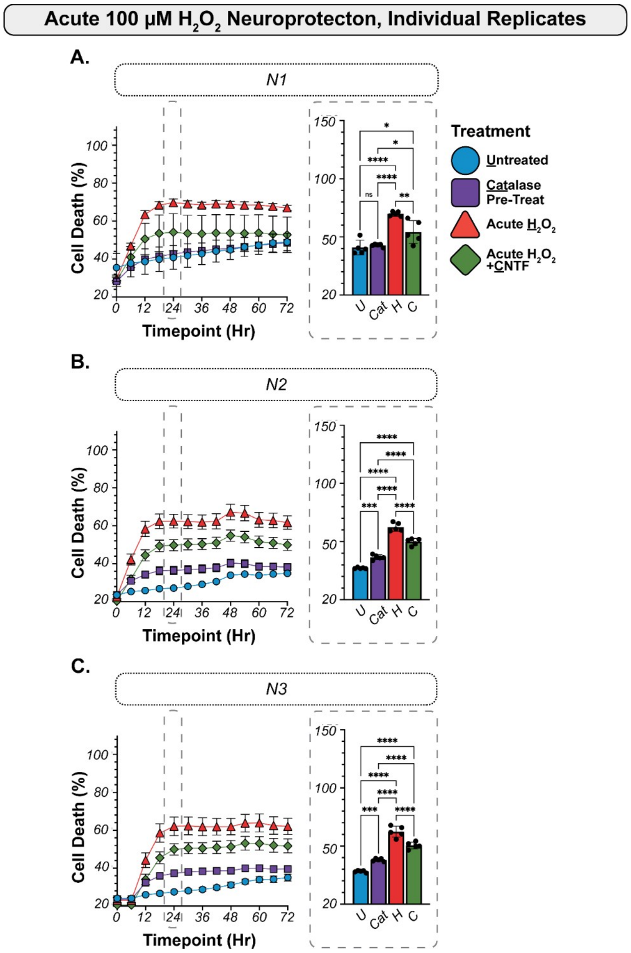


**Figure S15**: *Individual biological replicates for CNTF neuroprotection assay against acute 100 µM $H_2O_2$ exposure in MCN.*
(A - C) Three unique preparations of MCN used to evaluate neuroprotective potential of CNTF against acute (30 min.) 100 µM $H_2O_2$ with the measured time course and representative 24-hour time point with statistics. Statistics: Mixed model Two-Way ANOVA, multiple comparisons, with post-hoc Tukey correction test; data presented as mean ± SD; α = 0.05; ** p < 0.01; *** p < 0.001; **** p < 0.0001; ns = not significant; n = 5 (•).

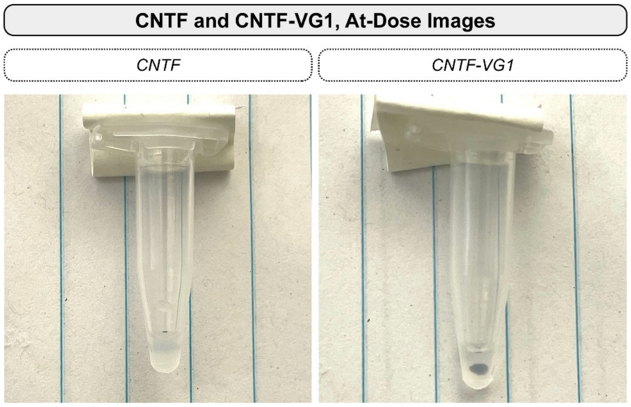


**Figure S16**: *CNTF and CNTF-VG1 turbidity images for at-dose concentrations (4 ºC).*
Qualitative CNTF (left; 2 mg mL$^{-1}$) and CNTF-VG1 (right; ~5 mg mL$^{-1}$) images after ~2 weeks at 4 ºC show heightened turbidity in CNTF compared to CNTF-VG1. A black dot was drawn behind the solutions to show relative degree of clarity.

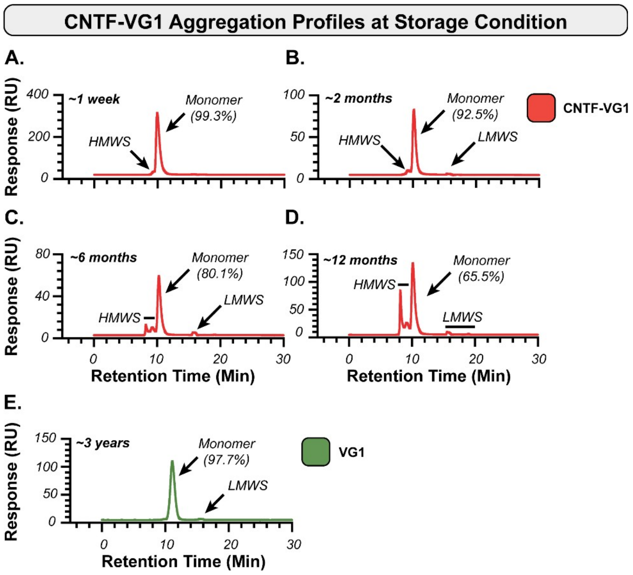


**Figure S17**: *CNTF and CNTF-VG1 aggregation over time under storage conditions.*
Representative SEC traces taken of CNTF-VG1 approximately (A) 1 week, (B) 2 months, (C) 6 months, and (D) 12 months after expression when stored as a liquid formulation at 4 ºC; shown with annotations for total monomeric content. (E) Representative SEC trace taken of a sample of VG1 after approximately 3 years of storage under identical conditions.

### ❖ SUPPLEMENTAL MATERIALS AND METHODS:

**H1-hESC Brn3B-mCherry⁺-derived retinal ganglion cell differentiation:** Human retinal ganglion cells (RGC) were differentiated from H1-hESC Brn3b-mCherry⁺ (ESC) as previously described[64]. In brief, ESCs were seeded (D: -1) on Matrigel- (354227, Corning) coated plates in fresh mTeSR™ Plus medium (100-0276, StemCell) as small clumps and then stepwise differentiated into neural rosettes. First, ESCs were treated with induction medium (DMEM/F12, 1% N2 Supplement, 2% B27 Supplement with 1 ng $mL^{-1}$ of Noggin (6057-NG, R&D Systems), Dkk-1 (5439-DK, R&D Systems), and IGF-1 (291-G1, R&D Systems)) for 72 hours (D: 0-3). Then, cultures were further driven to a retinal progenitor cell lineage with neural rosette differentiation medium (DMEM/F-12, 1% N2 Supplement, 2% B27 supplement; 10 ng $mL^{-1}$ of Noggin, Dkk-1, and IGF) for ~2.5 weeks (D: 4 ~ 21) until they formed robust embryoid bodies. Embryoid bodies were manually isolated (D: 18-21) using a sterile 21-G insulin syringe, re-seeded on Matrigel-coated plates and incubated for 72 hours (D: 0-2) in RGC Initiation Medium (RGC Basal medium (DMEM/F-12 : Neurobasal (1:1) medium, 0.05% N2 Supplement, 0.5% B27 Supplement, 5% L-Glutamine, 1% β-Mercaptoethanol, 1 µg $mL^{-1}$ bovine serum albumin (BSA), 1X ITS-G (41400045, ThermoFisher) with 250 ng $mL^{-1}$ Shh, 100 ng $mL^{-1}$ FGF8 (423-F8, R&D Systems), and 3 µM DAPT (D5942, Sigma)). Cultures were then treated with RGC Differentiation Medium (RGC basal medium with 250 ng $mL^{-1}$ Shh, 100 ng $mL^{-1}$ Follistatin, and 3 µM DAPT) for 72 hours (D3 - 5) followed by RGC maturation medium (RGC Basal Medium with 100 ng $mL^{-1}$ BDNF (248-BDB, R&D Systems), 5 ng $mL^{-1}$ NT4 (268-N4, R&D Systems), 10 ng $mL^{-1}$ CNTF, 10 µM Forskolin, 10 µM Y27632, 400 µM db cAMP, and 3 µM DAPT) for approximately 2 weeks (D: ~18). Mature RGC's were enriched through FACS sorting following Accutase (Corning, 25-058-CI) dissociation and cell filtering (Corning, 431750) wherein mCherry⁺ cells were isolated and seeded onto Matrigel-coated plates in fresh RGC Maturation Medium. Cells were left to grow neurite protrusions for ~1 week prior to any testing.

**JAK/STAT3 activation in H-hESC Brn3B-mCherry⁺ derived retinal ganglion cells quantification:** FACS-sorted H1-hESC Brn3B-mCherry⁺-derived RGC were gently washed with PBS (2X) and incubated overnight in CNTF-free RGC Maturation Medium (medium) to allow for clearance of any $pSTAT3^{Tyr705}$. The following day, cells were treated with either: 1) CNTF-free medium, 2) medium with 100 $mL^{-1}$ CNTF, and 3) medium with a molar equivalent of CNTF-VG1. After 30 minutes of exposure, cells were gently washed (3X) with PBS and cell lysate was collected with RIPA buffer (9806, Cell Signaling) with protease/phosphatase inhibitor, further dissociated through probe sonication and cleared of insoluble cell/tissue components via centrifugation (10 min; 4 ºC, >20,000 x g). Protein content was measured using a BCA assay (23225, ThermoFisher), and ~20 µg of tissue lysate was separated *via* electrophoresis, transferred to a nitrocellulose membrane, and blocked for 1 hour at room temperature with blocking buffer (37516, ThermoFisher). Staining and imagining were conducted in two stages: first, membranes were co-labeled with antibodies for $pSTAT3^{Tyr705}$ (1:1000, Rb; 9145, Cell Signaling) and β-Actin (1:10,000, Ms; 3700; Cell Signaling), washed, and subsequently stained with a corresponding secondary antibody cocktail (1:10,000, anti-Rb; LIC-926-3221; 1:20,000 anti-ms; LIC-926-68052, LiCOR) and imaged (LiCOR Odyssey). Membranes were stripped (1 hr., room temperature; 21059, ThermoFisher), washed, re-blocked and then incubated with antibodies for STAT3 (1:1000, Rb; 12640, Cell Signaling) and β-Actin, washed and subsequently stained with a corresponding secondary antibody cocktail and imaged (LiCOR Odyssey). For quantification, band intensities were measured in Fiji[43] (ImageJ 1.54 p) with the Gel Analyzer tool. Band intensities for $pSTAT3^{Tyr705}$ and STAT3 were normalized to β-Actin and quantified for each condition as a ratio of $pSTAT3^{Tyr705}$/STAT3 and reported.

**AF488 labeling:** Constructs were labeled with AF488 according to standard protocols. In brief, CNTF-VG1 was buffer-exchanged into 0.1 M sodium bicarbonate buffer (pH 8.3) using a desalting column (89882, ThermoFisher) and adjusted to a final concentration of 5 mg $mL^{-1}$. It was then mixed with AF488-NHS (A20000, ThermoFisher) and left to react in the dark for 4 hours at room temperature; free dye was removed with dye removal columns (2X; PI22858, ThermoFisher), buffer-exchanged into PBS and then dialyzed (3500 MWCO; PURD35050, Millipore Sigma) against PBS (4 ºC, 24 hours) to remove unreacted AF488-NHS. Samples of the unreacted protein, crude reaction, and purified sample were collected and run on SEC as previously described with the excipient being monitored in UV-Vis (λ = 280 nm, 495 nm) and fluorescence (λ = 495) to verify successful conjugation and free-dye removal. Final DoL (target: ~1.0 per molecule) was measured by UV-Vis spectrometry (Unchained Labs) using the relevant extinction coefficient and correction factor.

**Fluorescence correlation spectroscopy:** To evaluate the formation of CNTF-VG1-HA complexes, the diffusion time of Alexa 488-labeled CNTF-VG1 was monitored upon HA binding. Protein solutions (50 nM) were prepared in porcine vitreous, rat vitreous, neat PBS, and 1% 60 kDa HA in PBS. Data acquisition was performed using a Leica SP8 Laser Scanning Confocal Microscope equipped with a water immersion objective. Photon traces were recorded continuously for 27 minutes, with autocorrelation curves generated every 180 seconds. Data analysis was conducted using the Single Molecule Detection (SMD) FCS Wizard within the Leica LAS X software. To account for the two diffusing species (bound CNTF-VG1 and free CNT-VG1), the autocorrelation curves were fitted to a two-component model. The fitting parameters included the number of molecules (N), the fraction of molecules in triplet state (T), the triplet relaxation time ( $\tau_T$ ), the diffusion time ( $\tau_D$ ), and the structural parameter (s) representing the ratio of the radial to axial dimensions of the focal volume:

$$G(\tau) = \frac{1}{N}\left(1 + \frac{T}{1-T}e^{T/\tau_T}\right)\left(1 + \frac{\tau}{\tau_D}\right)^{-1}\left(1 + \frac{\tau}{s^2\tau_D}\right)^{-1/2}$$

An Alexa 488 standard dye solution with known diffusion time at room temperature was used to calibrate the lateral beam waist ($\omega_{xy}^2$). This calibration enabled the determination of the diffusion coefficient (D) for CNTF-VG1 across the various media using the following relationships:

$$D = \frac{\omega_{xy}^2}{4\tau_D}$$

$$V_{eff} = \pi^{3/2}\omega_{xy}^2 z_0$$

**Vitreous HA quantification:** Primary, pooled, vitreous samples for human, minipig (pig), cynomolgus monkeys (cyno) and rabbit were purchased from commercial suppliers (BioIVT, BioChemed). Rat (Sprague-Dawley) vitreous was isolated in-house. Frozen vitreous samples were thawed overnight at 4 ºC, centrifuged (>20,000 x g) to remove insoluble material, and the resulting supernatant was diluted (10X ~ 200X) in pre-chilled, sterile-filtered, phosphate buffered saline (PBS). Diluted samples (2 - 3 replicates) were measured for HA-content using an HA-ELISA kit (K-1200, Echelon Biosciences) according to manufacturer instructions with reference to an HA ladder.
